# Evolutionary history and microbial cross-feeding shape lifestyle-stratified dominance of *Bifidobacterium longum* subspecies in infants

**DOI:** 10.64898/2026.07.13.737801

**Authors:** Wei Guo, Wen Zhang, Lili Yang, Trine Zachariasen, Xuanji Li, Jakob Stokholm, Min Dai, Jonathan Thorsen, Søren J. Sørensen, Urvish Trivedi

## Abstract

*Bifidobacterium longum* (*Bl*.) is a key early-life gut symbiont, yet its evolutionary origin and mechanisms underlying the global biogeographic distribution of its subspecies remain poorly resolved. Here, we compiled a global genomic atlas of >7,000 MAGs/genomes from infants, domesticated animals, non-human primates, and ancient humans. High-resolution phylogenomic and functional analyses expanded infant-associated subspecies to five. Compared with non-human primates, *B. longum* was more prevalent in domesticated animals and ancient humans dating from 150 to 1,500 years ago. Moreover, human- and livestock-derived lineages from the same geographic regions clustered together, suggesting potential host-associated transmission. Ecologically, *Bl. infantis* and *Bl. longum* predominated in non-Western and Western infants, respectively, independent of breastfeeding, delivery mode, or antibiotic exposure. Instead, their distribution was associated with co-occurring microbes and HMO-driven cross-feeding interactions. These findings explain subspecies differentiation in the gut microbiota of Western and non-Western infants and provide a framework for community-mediated interventions in early life.

## Graphical Abstract

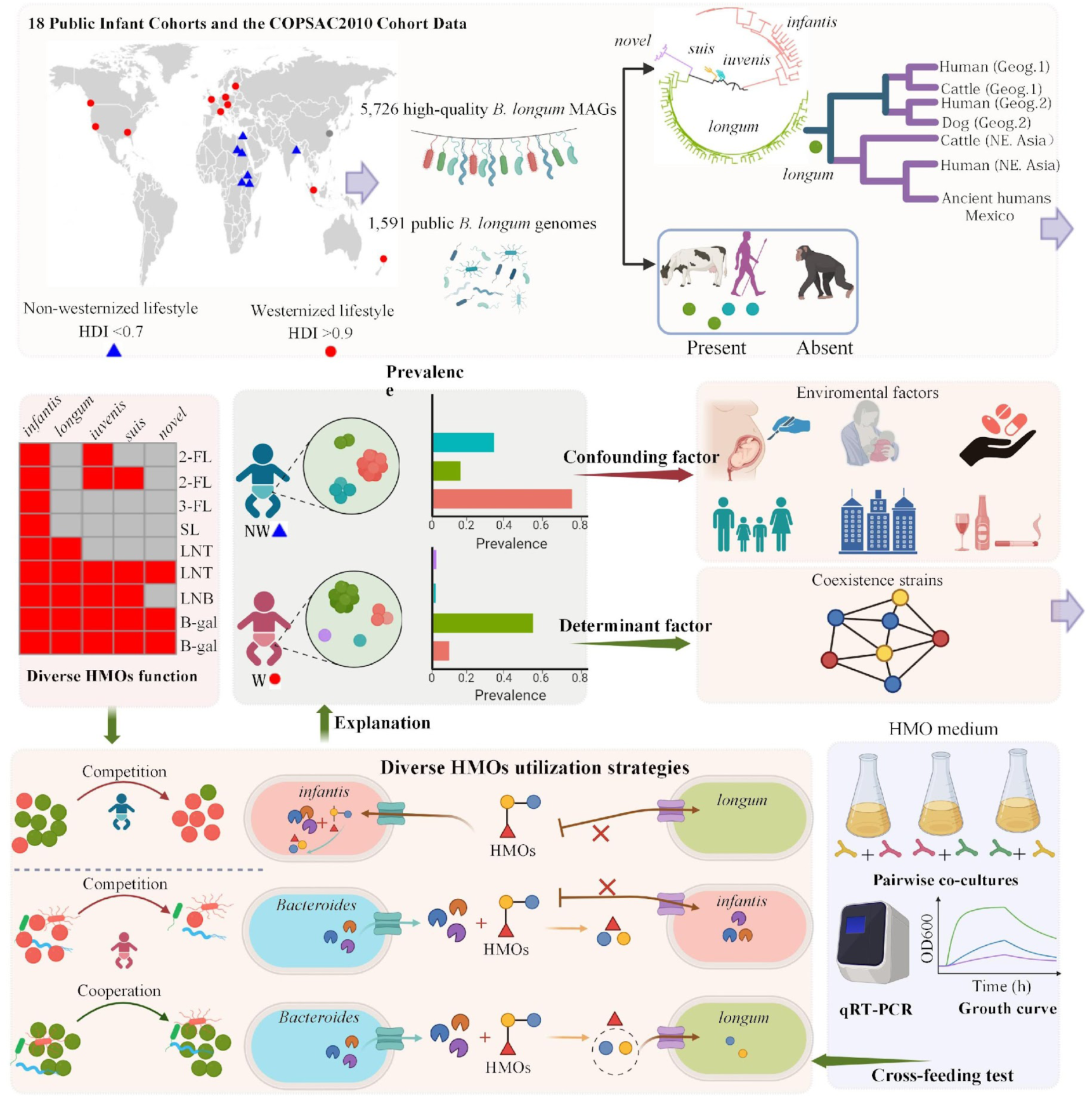

## Introduction

*Bifidobacterium longum* is among the earliest colonizers of the infant gut ^1–3^ and confers multiple health benefits during early development, including mitigation of dysbiosis ^4^, metabolism of human milk-derived carbohydrates ^5^, and promotion of immune maturation ^6^. For a long time, *B. longum* has been considered to comprise three subspecies ^3,7^. The Genome Taxonomy Database (GTDB) classified *Bl. infantis* as an independent species. Recently, large-scale genomic studies have reclassified human-associated *B. longum* strains into *B. infantis* and three *B. longum* subspecies: *longum*, *suis*, and *B. longum novel/*X ^5,8^. However, the genetic distance between *Bl. infantis* and some *B. longum* subspecies did not meet the criterion for species-level differentiation, as their average ANI divergence was less than 5%. Controversial, some studies have reported ambiguity surrounding the subspecies *Bl. iuvenis* ^9^, suggesting that it likely belong to *Bl. suis* or represented an independent subspecies. Consequently, it is still unclear whether infant-associated *B. longum* diversity follows a four-subspecies framework (*Bl*. *longum*, *Bl. infantis*, *Bl. suis*, and *Bl. novel*/X) or a five-subspecies framework (*Bl. longum*, *Bl. infantis*, *Bl. suis*, *Bl.iuvenis*, and *Bl. novel*/X). Beyond taxonomy, the evolutionary origin of *B. longum* remains unclear, largely due to the scarcity of cross-species evolutionary studies. Given that *Bl. suis* was mainly isolated from domesticated animals ^10^, it raises the possibility that certain some *B. longum* subspecies may also have arisen through host transmission associated with domestication.

Early-life gut microbiota assembly follows lifestyle-associated trajectories, reflecting distinct ecological selective pressures in non-Western versus Westernized populations ^11^. The differential prevalence of *B. longum* subspecies between Westernized and non-Western populations has long been attributed to differences in early-life environments, particularly breastfeeding practices, delivery mode, and antibiotic exposure ^12,13^. Despite these longstanding hypotheses, direct empirical evidence linking specific environmental exposures to *B. longum* subspecies-level population structure remains scarce.

Carbohydrate metabolism, particularly HMO utilization, is central to *Bifidobacterial* fitness in the infant gut and has been proposed as a key determinant of *B. longum* dominance across populations ^1,6,13^. *Bl*. *infantis* and *Bl. iuvenis* possess extensive genetic repertoires for direct HMO uptake and intracellular degradation, whereas *Bl*. *longum* generally lacks complete pathways for utilizing HMOs such as 2’-fucosyllactose (2’-FL), 3’-fucosyllactose (3’-FL), 3’**-**sialyllactose (3’-SL), and 6’**-**sialyllactose (6’-SL), and is thought to rely on metabolic byproducts generated by other HMO- degrading microbes ^3^. These genomic disparities point to distinct ecological strategies, potentially involving niche partitioning, competition, or cross-feeding interactions. However, the subspecies-specific interaction modes of *B. longum* within the infant gut ecosystem remain to be systematically characterized.

Taken together, a comprehensive analysis within a subspecies-level taxonomy framework is needed to accurately assess *B. longum* distribution during early life across distinct lifestyles. More critically, the mechanisms by which Western and non-Western infant gut environments shape lifestyle-stratified dominance of subspecies and functional strategies within *B. longum*, particularly through the interplay of environmental pressures, microbial metabolism, and inter-microbial interactions remain largely unexplored. Addressing these knowledge gaps is essential for advancing our understanding of infant gut microbiome assembly and the global divergence of early-life microbial communities.

In this study, we performed large-scale metagenomic assembly to reconstruct over 5,700 high-quality *B. longum* genomes from the Copenhagen Perspective Studies on Asthma in Childhood (COPSAC_2010_) cohort and publicly available metagenomes spanning 18 cohorts and diverse lifestyles. Comparative genomic and distribution analyses of *Bl. longum*, *Bl. iuvenis* and *Bl. suis* across multiple host sources were used to clarify the evolutionary relationships and origins of the human-associated lineage. The unique pangenomes of each *B. longum* subspecies were constructed to resolve the global distribution of subspecies-level lineages in early infancy. We further characterized the HMO metabolic potential of the subspecies and key co-occurring species through comparative genomics, statistics, and *in vitro* cultivation. Together, these analyses provide an unprecedented, integrated view of how *B. longum* subspecies adapt to divergent early-life environments across the globe.

## Results

### The Distribution of Five Distinct *B. longum* Subspecies in Infants

A total of 5,726 high-quality metagenome-assembled genomes (MAGs) of *B. longum* (>95% completeness, <5% contamination) were reconstructed from 6,674 shotgun metagenomic sequenced samples from 19 cohorts (Table S1), including the newly released COPSAC_2010_ data from 1-month-old infants.

Phylogenetic reconstruction resolved *B. longum* genomes into five evolutionarily divergent clades, and represent *Bl. infantis*, *Bl. longum*, *Bl. iuvenis*, *Bl. suis*, and *Bl. novel* (Fig. 1A). Pairwise average nucleotide identity (ANI) analysis of the genomes showed substantially lower intra-subspecies distances (<2%) than inter-subspecies distances (2–5.5%) (Fig. 1B). Within *Bl. novel*, ANI divergence was especially low (<1%), while the smallest inter-subspecies distance was observed between *suis* and *iuvenis* (2.04%). Among these lineages, only *Bl. infantis* exhibited an average ANI distance greater than 5.0% relative to both *Bl. longum* (5.03%) and *Bl. novel* (5.31%). In bacterial taxonomy, genomes sharing >95% ANI are generally considered members of the same species ^14^, whereas lineages displaying appreciable genomic divergence within this range are often recognized as subspecies-level taxa. Therefore, defining *Bl. infantis* as a fully distinct species separate from *B. longum* may be taxonomically debatable. Infant cohorts in most westernized countries exhibited four *B. longum* clades at least (Fig. S1A; Table S2), whereas those in non-westernized regions typically harbored three clades (Fig. S1B; Table S2), a pattern consistent with that observed in Chinese populations (Fig. S1C; Table S2). Among them, *Bl. infantis* and *Bl. longum* were ubiquitous across countries, whereas the *Bl. novel* subspecies was largely restricted to westernized populations. *Bl. iuvenis* was detected in most countries, while *Bl. suis* showed a limited geographic distribution, being recovered only in Denmark, Sweden, New Zealand and Zimbabwe. Notably, all five subspecies were recovered exclusively in the COPSAC_2010_ cohort. *Bl. longum* exhibited no clear separation by lifestyle groups (western vs non-western) (Fig. 1C). In contrast, *Bl. infantis*, *Bl. iuvenis*, and *Bl. suis* showed strict lifestyle-based separation (Fig. 1C). Among them, *Bl. infantis* forms three distinct clades: a Western lifestyle–associated clade, a clade predominantly comprising MAGs from Bangladesh and Kenya, and a third clade mainly consisting of MAGs from Zimbabwe, South Africa, and Mozambique. In contrast, *Bl. iuvenis* and *Bl. suis* cluster into two clearly separated branches corresponding to Western and non-Western lifestyles. These findings were corroborated by Principal Coordinate Analysis of core gene presence/absence across all MAGs (Fig. S1 D-G), where *Bl. infantis*, *Bl. iuvenis*, and *Bl. suis* again demonstrated strict lifestyle segregation (Fig. S1 E-G).

**Fig. 1.**
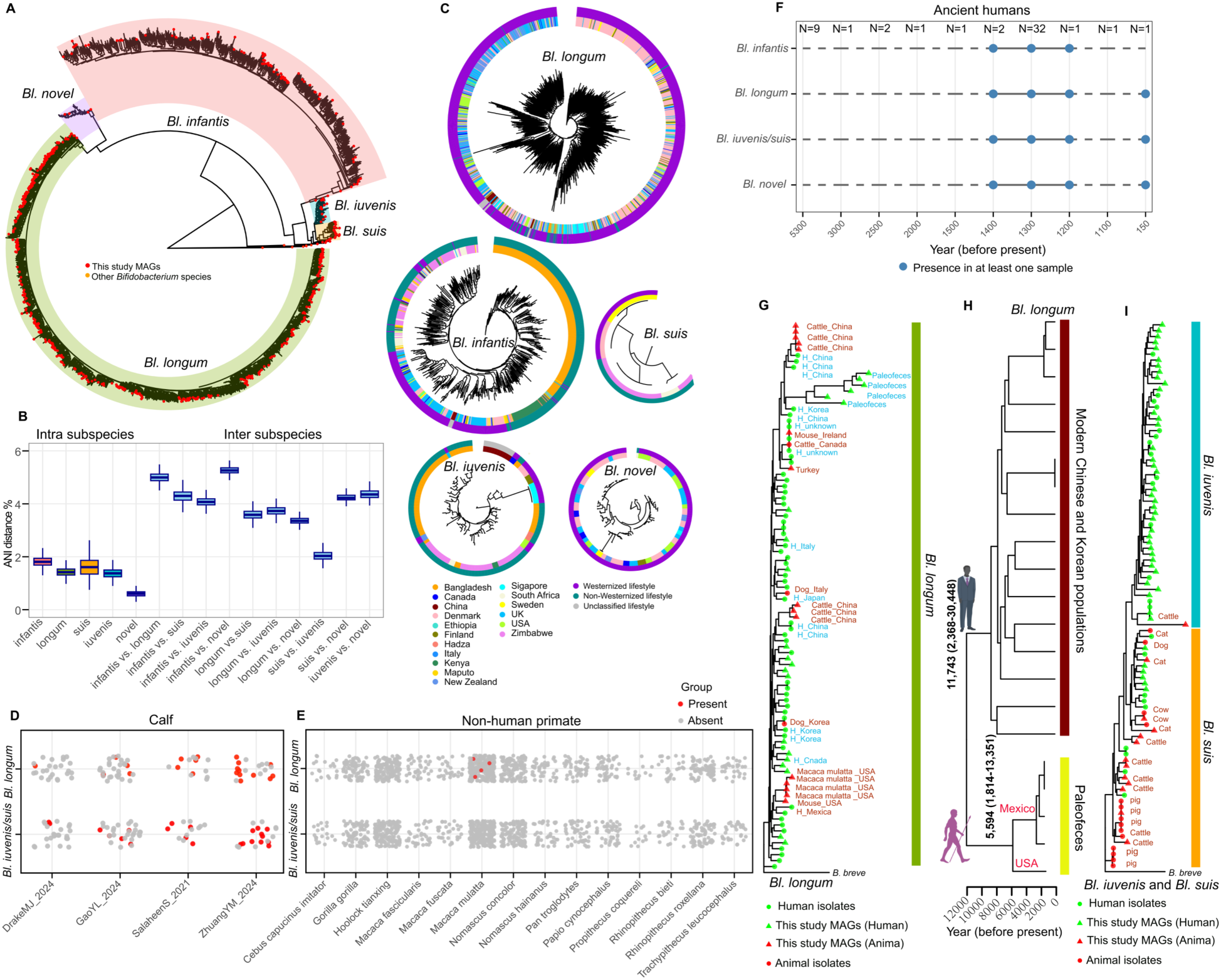
The Five Distinct Subspecies of *B. longum* in Early Life. **(A)** Whole-genome phylogenetic tree of representative subsets from the five *B. longum* (*Bl*) subspecies; **(B)** Genetic distances within subspecies (intra-subspecies) and between clades (inter-subspecies) are shown as pairwise average nucleotide identity (ANI) distances; **(C)** Phylogenetic representation of core genomes in *Bl. longum*, *Bl. infantis*, *Bl. iuvenis*, *Bl. suis*, and *Bl. novel*; **(D)** Prevalence patterns of *B. longum* subspecies across calf metagenomes derived from five independent studies; **(E)** Prevalence patterns of *B. longum* subspecies across metagenomes from 14 non-human primate species; **(F)** *B. longum* subspecies detected in ancient human metagenomes; **(G)** Phylogenetic tree of *Bl. longum* derived from diverse host sources, including ancient human samples (∼1,300 years before present); **(H)** Time-calibrated phylogeny of modern Asian and ancient American *B. longum* strains inferred using a Bayesian molecular clock. Numbers at internal nodes indicate median divergence time estimates, and node bars represent the corresponding 95% highest posterior density (HPD) intervals; **(I)** Phylogenetic tree of *Bl. iuvenis* and *Bl. suis* derived from diverse host sources.

To investigate the potential origin of *B. longum*, we assessed the prevalence of the *B. longum* in calves, non-human primates, and palaeofaeces. *Bl. iuvenis/suis* and *Bl. longum* were prevalent in calves (Fig. 1D). In contrast, *B. longum* was largely absent in 37 non-human primates, with only 4 Macaca mulatta out of 934 non-human primates’ samples harboring the *Bl. longum* subspecies (Fig. 1E). No *B. longum* subspecies were detected in palaeofaeces dated to 1,500–5,300 years before present (Fig. 1F). In contrast, *Bl. infantis*, *Bl. longum*, *Bl. iuvenis/suis*, and *Bl. novel* were all detectable in samples dated to 1,200–1,400 years, whereas only *Bl. longum*, *Bl. iuvenis/suis*, and *Bl. novel* were recovered from more recent samples (∼150 years before present) (Fig. 1F). These observations do not directly support a primate-associated evolutionary history for *B. longum* but instead suggest a potential link transition between humans and domesticated animals. We therefore further examined the evolutionary relationships of *Bl. longum*, *Bl. iuvenis* and *Bl. suis* subspecies across different hosts. The MAGs and reference genomes used for phylogenetic tree construction are listed in Table S3. For *Bl*. *longum*, the subspecies recovered from Native American individuals dating to ∼1,300 years before present clustered with those from human populations in China and Korea, as well as cattle from China, while showing a more distant relationship to *longum* from Macaca mulatta (Fig. 1G). In addition, *Bl. longum* from Chinese cattle consistently clustered with human isolates from the Chinese population, and those from Korean dogs consistently clustered with human isolates from the Korean population (Fig. 1G). Bayesian molecular clock analysis estimated the divergence time between the modern Asian *Bl. longum* strains and the ancient American strains (Mexico and USA combined) to have a tMRCA of 11,743 years (95% HPD: 2,368–30,448 years; Fig. 1H). For *Bl. suis*, human-derived strains clustered with those from domesticated animals, including cattle, cats and dogs, and formed a distinct lineage separate from *Bl. iuvenis*. Notably, the only animal-derived *Bl. iuvenis* MAG clustered with *Bl. iuvenis* subspecies of Chinese origin (Fig. 1I). This suggests potential host-associated transition or ongoing microbial exchange between human and domesticated animals.

### Divergent Carbohydrate Metabolism Repertoires Across *B. longum* Subspecies

Hierarchical clustering of the heatmap based on the overall distances of the Carbohydrate-Active enZymes (CAZymes) functional profiles revealed considerable functional diversity among the five *B. longum* subspecies, with *Bl. infantis* being the most dissimilar compared with other subspecies based on the overall distance of the CAZymes functional profiles (Fig. S2A). The distinct clustering observed in NMDS (Non-Metric Multidimensional Scaling) analyses based on all CAZy profiles (Fig. 2A), HMO (Fig. S2B), host glycans (Fig. S2C), diet glycans (Fig. S2D), and starch (Fig. S2E) gene content reveals functional differentiation among subspecies, suggesting heterogeneity in carbohydrate metabolism. Notably, the *Bl. iuvenis* and *Bl. suis* exhibit highly similar profiles across all five gene content categories and are seen to consistently cluster together when doing distance analysis (Jaccard distance). Based on the annotation results of CAZyme genes, *Bl. longum* contains the significantly greatest number of CAZyme genes (*p* < 0.05 with each subspecies), whereas *Bl. infantis* contains the lowest (Fig. 2B). However, the significantly highest number of HMO degradation genes was observed in *Bl. infantis* (*p* < 0.05 with each subspecies), followed by *Bl. iuvenis*, *Bl. suis*, and *Bl. longum*, with *Bl. novel* possessing the fewest (Fig. 2B). Furthermore, *Bl. longum* and *Bl. iuvenis* possessed the most non-HMO glycan-degrading genes, followed by *Bl. novel* and *Bl. suis*, with *Bl. infantis* having the fewest (Fig. 2B). Interestingly, the *Bl. novel* subspecies harbors a significantly greater number of genes involved in starch metabolism (*p* < 0.05 with each subspecies) (Fig. 2B).

**Fig. 2.**
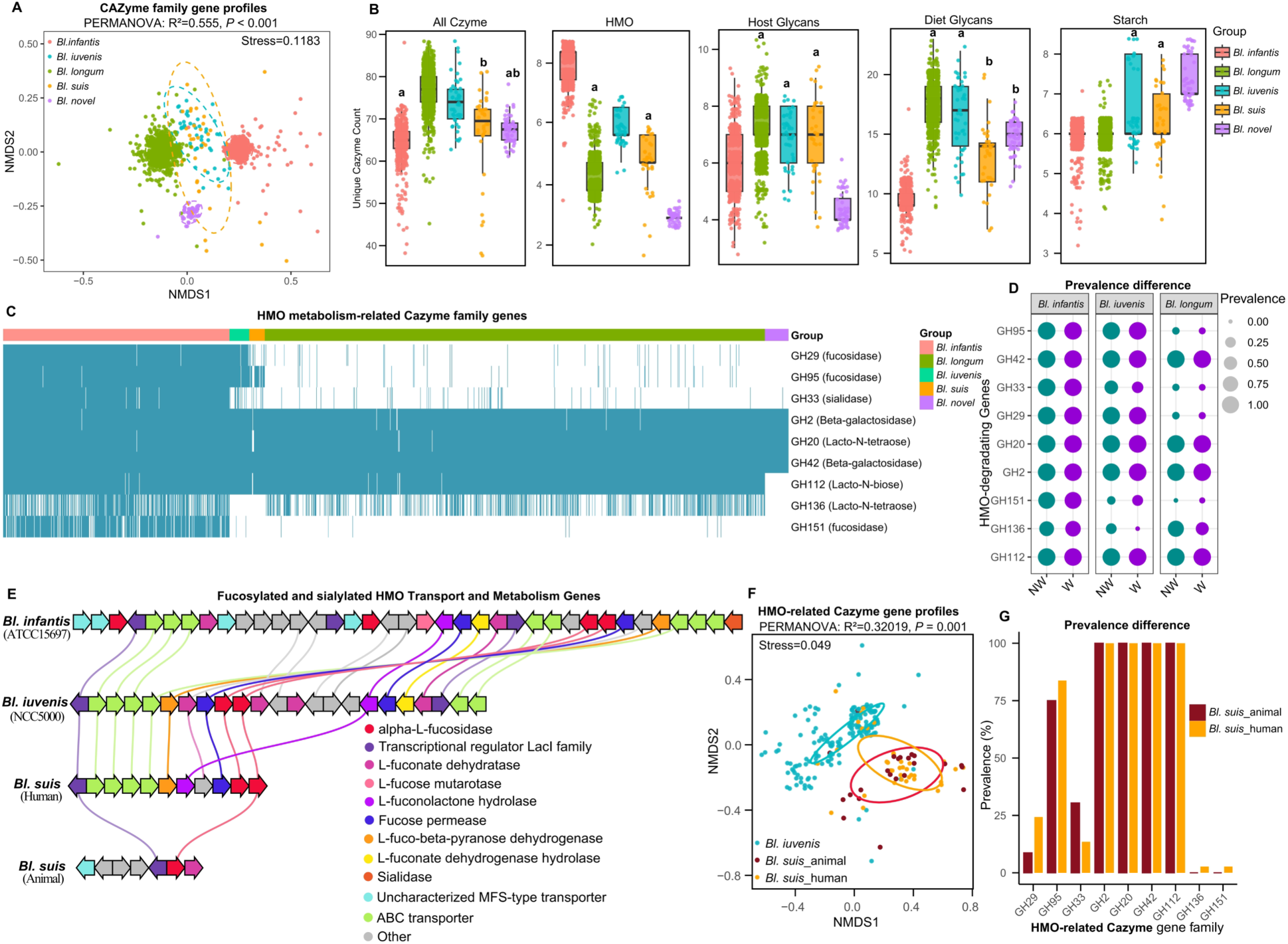
Functional Diversity of *B. longum* Subspecies. **(A)** Non-Metric Multidimensional scaling (NMDS) ordination of CAZy families in each genome showing distinct inter- and intra-subspecies clustering within *B. longum* subspecies using Jaccard distances; **(B)** Count of unique all CAZyme, HMO-digesting, glycans-digesting, and starch-digesting genes across the five *B. longum* subspecies; **(C)** Presence and absence of key HMO-degrading genes differ significantly across the five *B. longum* subspecies MAGs and genomes; **(D)** Comparative prevalence of HMO utilization genes in *Bl. infantis*, *Bl. longum*, and *Bl. iuvenis* across Western and non-Western lifestyles; **(E)** Conserved synteny of fucosylated HMO-degrading genes in representative genomes of five *B. longum* subspecies; **(F)** NMDS using Bray-Curtis distances of CAZy families associated with HMO utilization among *Bl. iuvenis*, human-derived *Bl. suis,* and animal-derived *Bl. suis*; **(G)** Prevalence of HMO utilization–associated genes (GH) in human- and animal-derived *Bl. suis*. In the Kruskal–Wallis test, groups sharing the same letter (e.g., “a” and “a”) are not significantly different, indicating that the *p*-value is greater than 0.05.

A total of 26 glycoside hydrolase (GH) genes are broadly conserved across the five subspecies, predominantly involved in the degradation of host-derived and diet-derived glycans (Fig. S2F). The metabolic potential for HMO utilization varied significantly across the five subspecies. In detail, all subspecies possess the ability to metabolize lactose via beta-galactosidases (GH2 and GH42) (Fig. 2C). However, Lacto-N-tetraose (degraded by GH20) was the only HMO metabolized by all five subspecies. Specifically, *Bl. infantis* was capable of utilizing all HMO. The *Bl. iuvenis* and *Bl. suis* can both utilize fucosylated HMO, lacto-N-tetraose HMO, and lacto-N-biose HMO. Compared with *Bl. suis*, *Bl. iuvenis* possesses the GH29 α-fucosidase gene involved in the utilization of fucosylated HMOs, whereas this gene is absent in *Bl. suis* (Fig. 2C). *Bl. longum* utilized only Lacto-N-tetraose and Lacto-N-biose. In contrast, the *Bl. novel* was only able to metabolize Lacto-N-tetraose (Fig. 2C). Considerable intra-subspecies variation was observed in the distribution of GH33 and GH136 (Table S4). Specifically, only 46.15% of *Bl. iuvenis* MAGs encoded GH33, while GH136 was detected in 64.18% of *Bl. infantis* and 43.83% of *Bl. longum* MAGs, indicating substantial heterogeneity in HMO-utilization potential within subspecies. In terms of diet-derived glycans, the *Bl. infantis* is deficient in several relevant genes, including those required for xylan degradation (Fig. S2F). By contrast, gene absence in *Bl. novel* predominantly affects pathways related to host glycan and HMO metabolism. Notably, *Bl. novel* consistently harbors genes associated with starch utilization, including those encoding starch endo-α-1,6-glucosidase (GH176) and GH13_14 enzymes (Fig. S2F). Lifestyle shapes the prevalence of some HMO-related genes in *B. longum* (Fig. 2D). Specifically, GH136 in the *Bl*. *infantis* increases from 62% in non-Western populations to 71% in Western populations (Chi-square test, *p*<0.001), whereas it declines from 79% to 42% in *Bl*. *longum* across the same comparison (Chi-square test, *p*<0.001). *Bl. infantis*, *Bl. iuvenis*, human-derived *Bl. suis*, and animal-derived *Bl. suis* all harbor genes encoding α-L-fucosidases, except for animal-derived *Bl. suis*, they form a highly similar specialized gene cluster for the utilization of fucosylated HMOs (Fig. 2E). Further NMDS analysis based on the presence/absence of HMO utilization genes revealed that human-derived *Bl. suis* exhibits a bifurcated clustering pattern: a subset cluster with *Bl. iuvenis*, while another subset clusters with animal-derived *Bl. suis* (Fig. 2F). In contrast, animal-derived *Bl. suis* forms a distinct cluster, clearly separated from *Bl. iuvenis*. In terms of gene prevalence, GH29 and GH93 are significantly more prevalent in human-derived *Bl. suis* than in animal-derived *Bl. suis*, whereas GH33 shows the opposite pattern, with lower prevalence in human-derived *Bl. suis* (Fig. 2G).

### Lifestyle-associated Differences in *B. longum* Subspecies Distribution in Infants

Due to the high similarity between *Bl*. *iuvenis* and *Bl. suis* and the limited number of human-derived *Bl. suis* genomes, only ∼10 genes satisfied the criteria for designation as subspecies-specific genes in the *B. longum* pangenome. Consequently, *Bl*. *iuvenis* and *Bl. suis* were merged as an *Bl*. *iuvenis*/*suis* complex for the identification of unique pangenome genes. This yielded 148, 90, 74, and 135 marker genes for *Bl*. *infantis*, *Bl*. *longum*, *Bl*. *iuvenis/suis*, and *Bl*. *novel*, respectively (Table S5).

*Bl. infantis* (57.11%) and *Bl*. *iuvenis*/*suis* (32.71%) constituted substantially higher proportions in non-Westernized countries than in Westernized countries (*Bl*. *infantis*: 10.04%; *Bl*. *iuvenis*/*suis*: 0.75%; Fig. 3A; Fig. S3A; Table S6). The proportions of *Bl. infantis* in Kenya populations were nearly ubiquitous (99.05%), in contrast some Western countries were almost lacking it (New Zealand: 1.86%; Finland: 4.12%; Table S6). The same pattern was seen for *Bl. iuvenis/suis,* which was prevalent in non-Westernized infants, but rarely seen in Western-style infants. In most Western-style countries, its proportion was less than 1% (Fig. 3A). Conversely, *Bl. longum* demonstrated significantly higher prevalence in Western infants (53.88%), whereas its prevalence was lower in non-Western infants (17.03%). *Bl. novel* was detected in most of Western cohorts but exhibited relatively low prevalence (0% to 4.24% across countries; Fig. 3A). It was absent in non-Western populations, with the exception of South Africa, where it was detected in only one sample. In non-Westernized infants, the prevalence rates of *Bl. infantis* and *Bl. iuvenis/suis* rose after birth, peaking at around 6–12 months and 10–16 months of age, respectively (Fig. 3B). Subsequently, both declined with advancing age. In contrast, the prevalence rate of *Bl. longum* showed a slight decrease after birth before increasing again after 10 months of age (Fig. 3B). While the overall prevalence of *Bl. infantis* was lower in Westernized infants, its temporal trajectory mirrored that observed in non-Westernized populations (Fig. 3C). The prevalence of *Bl*. *longum* increased after birth and stabilized around 1 year of age. In contrast, *Bl. iuvenis*/*suis* and *Bl. novel* exhibited low prevalence rates across all ages. Notably, *Bl. novel* was only detected in infants under one year of age. Significantly higher prevalence (Mann-Whitney U test, *p*<0.05) of *Bl. infantis* and *Bl. iuvenis/suis* was observed among infants under 2 months, 3–12 months, and over 12 months in non-Westernized versus Westernized countries (Fig. S3 B). Conversely, the prevalence of *Bl. longum* and *Bl. novel* were significantly higher (Mann-Whitney U test, *p*<0.001) in Westernized countries (Fig. S3 B).

**Fig. 3.**
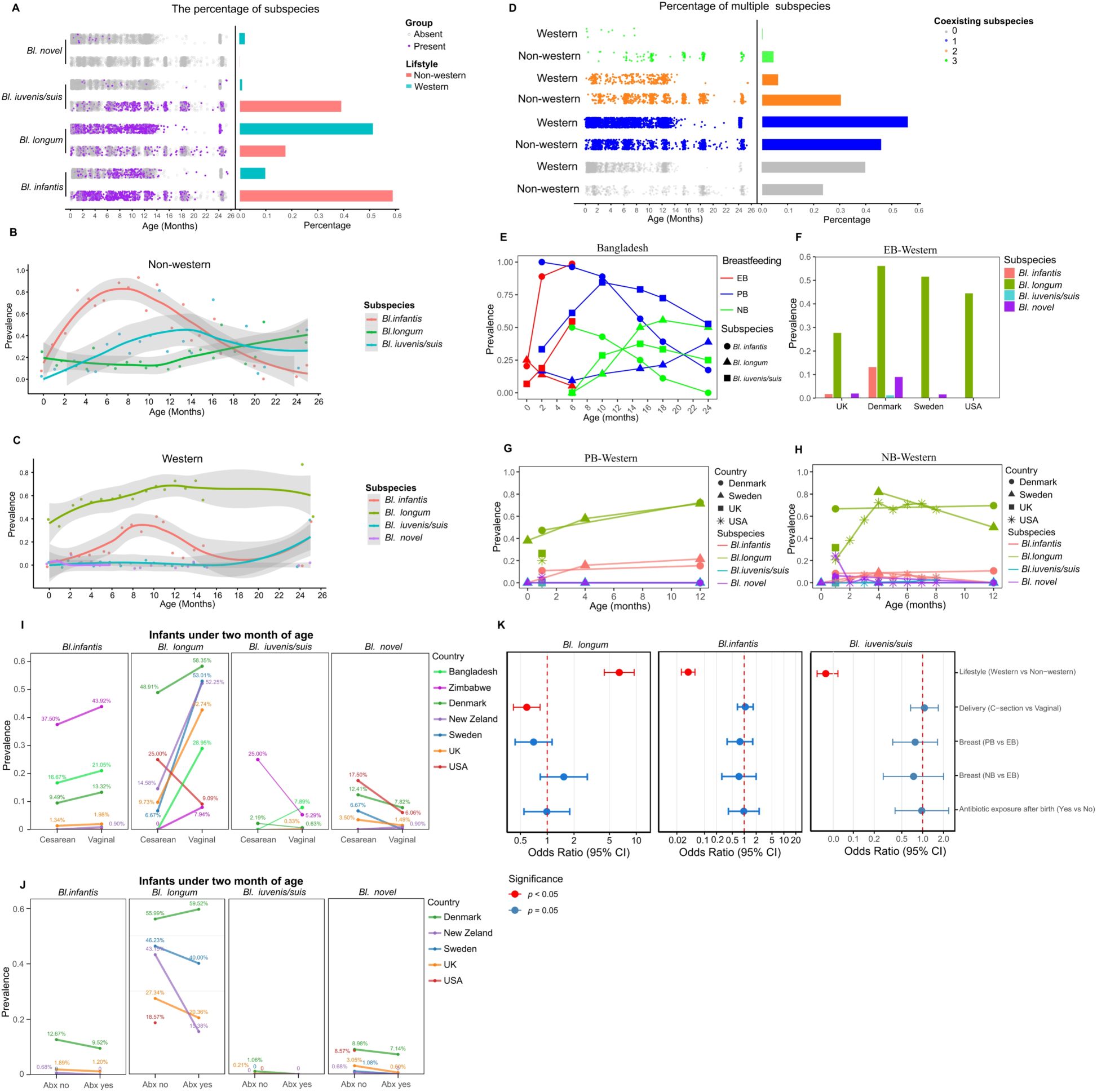
Prevalence patterns of *B. longum* subspecies in Western versus non-Western infants, and the impact of breastfeeding, delivery mode, and antibiotic exposure. (**A**) Percentage of present of distinct *B. longum* subspecies in infants with western and non-western lifestyles; The prevalence of distinct *B. longum* subspecies in infants with non-western **(B)** and western **(C)** lifestyles; **(D)** Percentage of individuals harboring multiple *B. longum* subspecies in infants with non-western and western lifestyles; **(E)** Prevalence dynamics of *B. longum* subspecies in Bangladesh infants by breastfeeding pattern (EB=Exclusive breastfed, PB=Partially breastfed, NB=Non-breastfed); Prevalence dynamics of *B. longum* subspecies in western lifestyle infants with exclusive breastfeeding **(F)**, partial breastfeeding **(G),** and no breast feeding **(H); (I)** Prevalence of *B. longum* subspecies in infants born by cesarean versus vaginal delivery; **(J)** Prevalence of distinct *B. longum* subspecies in infants with versus without intrapartum antibiotic exposure. **(K)** Factors associated with *B. longum* subspecies presence: results from generalized linear models. Abx: Antibiotic exposure.

Samples from non-Western populations showed a higher propensity for containing multiple *B*. *longum* subspecies, whereas Western populations more frequently exhibited no detectable subspecies (Fig.3D). Among infants with non-Western lifestyles, *Bl*. *infantis* demonstrated a marked predominance in relative abundance (Fig. S3C). *Bl*. *longum* showed markedly higher abundance in Western infants (Fig. S3D). However, *Bl*. *infantis* was also present at a considerable level of abundance in a small proportion of individuals. The abundances of *Bl. infantis* and *Bl. iuvenis/suis* were significantly higher (Mann-Whitney U test, *p*<0.001) in non-Western infants across all age groups (<2 months, 3–12 months, >12 months; Fig. S3 E), whereas *Bl. longum* and *Bl. novel* were significantly more abundant in Western infants (Fig. S3 E, Mann-Whitney U test, *p*<0.001).

We further assessed the influence of breastfeeding patterns, delivery mode, and antibiotic exposure on *B. longum* subspecies prevalence. Among Bangladesh infants, the prevalence of *Bl. infantis* and *Bl. iuvenis/suis* increased rapidly and significantly during exclusive breastfeeding (EB, from birth to 6 months old) (*Bl*. *infantis*: 20.45% to 97.14%; *Bl*. *iuvenis/susi*: 6.82% to 55.24%, Chi-square test, *p*<0.001), whereas *Bl. longum* prevalence decreased significantly (25.0% to 5.26%, Chi-square test, *p*=0.015) (Fig. 3E). In partially breastfed (PB) and non-breastfed (NB) infants, *Bl. infantis* prevalence gradually declined, *Bl. iuvenis/suis* exhibited a parabolic-like trend (initially rising then falling), and *Bl. longum* showed a steady upward trend (Fig. 3E). Collectively, *Bl. longum* demonstrated an inverse prevalence pattern relative to *Bl. infantis*. The abundance of *Bl. infantis* was significantly higher in the gut of exclusively and partially breastfed infants compared to formula-fed infants at time of sampling (Mann-Whitney U test, *p*<0.001). Furthermore, its abundance began to decline after two months of age (Fig. S3F). In contrast, no significant differences were observed in the abundance of *Bl. longum* and *Bl. iuvenis/suis* among infants under different feeding regimens (Mann-Whitney U test, *p*> 0.05). In the Western cohort, *Bl. longum* exhibited a high prevalence from 1 to 12 months of age across the EB (Fig. 3F), PB (Fig. 3G), and NB (Fig. 3H) patterns. In contrast, the prevalence of *Bl. infantis*, *Bl. iuvenis/suis* and *Bl. novel* remained low and stable from 1 to 12 months of age, with no significant variation across the different breastfeeding patterns. In western lifestyle, a higher relative abundance of *Bl. longum* and a lower abundance of *Bl. infantis* were observed in infants across all three breastfeeding patterns (Fig. S3 G, H, I), with the exception of PB Swedish infants at 4 months of age. Notably, the prevalence of *Bl. infantis* in non-Western infants delivered by cesarean section exceeded that in Western infants, regardless of delivery mode (Fig. 3I). Certainly, the prevalence of *Bl. infantis* and *Bl. longum* (except in the USA) were lower in cesarean-delivered infants than in vaginally delivered infants before two months of age. Within the same delivery mode, Western infants had a higher prevalence of *Bl. longum* than non-Western infants; notably, even Danish cesarean-born infants showed a higher prevalence than their non-Western, vaginally-born counterparts (Fig. 3I). Conversely, the prevalence of *Bl. novel* was higher following cesarean delivery (Fig. 3I). However, cesarean delivery did not consistently reduce the prevalence of *Bl. iuvenis/suis*; for example, prevalence increases in Bangladesh but decreases in Denmark and Zimbabwe (Fig. 3I). Intrapartum antibiotic exposure was associated with reduced prevalence of all subspecies in infants under two months of age compared to unexposed infants (Fig. 3J). However, in the COPSAC_2010_ cohort, no significant reduction in the prevalence of *Bl. longum* was observed following intrapartum antibiotic exposure (Chi-square test, *p* > 0.051). Using generalized linear models, Western lifestyle was the most significant factor affecting the prevalence of *Bl*. *infantis*, *Bl*. *longum*, *Bl*. *iuvenis/suis*, with a negative association for *Bl*. *infantis* and *Bl*. *iuvenis/suis* (all OR < 1, *p* < 0.001, logistic regression models) but a positive association for *Bl*. *longum* (all OR > 1, *p* < 0.001, logistic regression models) (Fig. 3K).

### Colonization Persistence of *Bl. infantis* and *Bl. longum* in Infants from Western and Non-Western Populations

Persistence analysis revealed distinct colonization patterns: *Bl. longum* persisted in Western but not in non-Western infants (Fig. 4A), *Bl. infantis* persisted exclusively in non-Western infants, *Bl. iuvenis/suis* only in Bangladeshi infants, and *Bl. novel* in none. Apart from *B. longum*, other genera such as *Bacteroides*, *Phocaeicola* and *Parabacteroides* showed strong persistence exclusively in Western infants. Within these other genera, only *B. breve* and *B. bifidum* maintained robust persistence in non-Western infants. Under a Westernized lifestyle, persistent positive correlation with multiple bacterial species was seen, including *Phocaeicola vulgatus*, *Parabacteroides distasonis*, *Phocaeicola dorei*, *Bacteroides fragilis*, *Bacteroides uniformis*, *B. bifidum*, *B. adolescentis*, and *B. pseudocatenulatum* (Fig. S4A, left). Under a non-Westernized lifestyle, *Bl. infantis* showed persistent positive correlations with species such as *B. bifidum* and *B. breve*, but persistent negative correlations with 11 different species including *B. pseudocatenulatum*, *Parabacteroides distasonis*, *Phocaeicola vulgatus*, and *Bacteroides ovatus* (Fig. S4A, right). Persistent species in Western infants carried significantly more genes involved in HMO metabolism than non-persistent strains (Fig. S4B, Mann-Whitney U test, *p*< 0.01). In contrast, neither the number nor the diversity of HMO metabolic genes differed significantly between persistent and non-persistent strains in non-Western infants (Fig. S4C, Mann-Whitney U test, *p*> 0.05). Among all HMO metabolism-related genes, the largest disparities of persistent MAGs vs non-persistent MAGs in Westernized infants were observed for genes encoding metabolism of 2-FL (GH95), 3-FL (GH29), and 3-SL (GH33) (Fig. 4B), which are notably absent in *Bl. longum*. In contrast, these differences were attenuated in non-Westernized infants (Fig. 4C). Statistically, in infants from Westernized lifestyles, the copy numbers of these genes were significantly higher in persistent strains than in non-persistent strains, whereas no significant difference was observed in non-Westernized infants (Fig. 4D). Similarly, in Westernized infants, the copy numbers of GH29, GH95, and GH33 were positively correlated with species persistence percentage (Fig. 4E), whereas no such correlation was observed in non-Westernized infants (Fig. 4F). Further, in Western countries, MAGs carrying GH29, GH95, and GH33 among persistent colonizers were primarily from genera such as *Bacteroides*, *Phocaeicola* and *Parabacteroides* (Fig. S4D), whereas in non-Western countries, such MAGs were mainly derived from the *Bifidobacterium* genus (Fig. S4D). Specifically, all species belonging to *Bacteroides*, *Phocaeicola*, and *Parabacteroides* possessed GH29, GH95, and GH33 but were devoid of GH112 (Fig. 4G), which was opposite the pattern seen in *Bl. longum*.

**Fig. 4.**
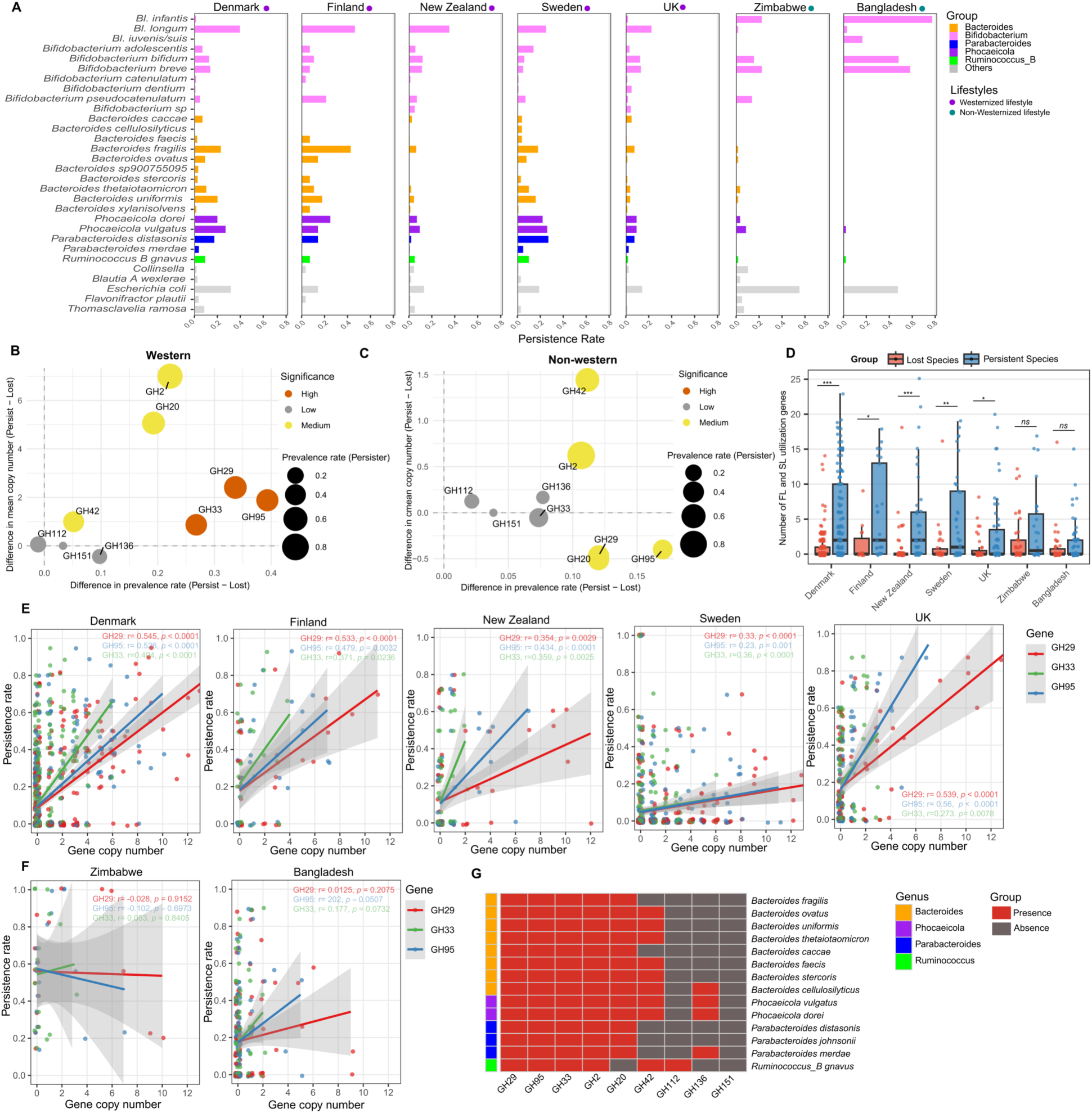
Characteristics of persistent and non-persistent gut species from Western and non-Western infants. **(A)** Species persistence rates in infants from non-Westernized (Bangladesh, Zimbabwe) and Westernized (Denmark, Sweden, Finland, New Zealand, UK) populations during the first year of life; Differences in prevalence rate and mean copy number of HMO utilization genes between persistent and lost species in Western **(B)** and Non-western **(C)** infant gut microbiomes; High: prevalence rate difference > 0.25 and mean copy number difference > 1; Medium: not meeting High criteria, but either rate difference > 0.1 or copy number difference > 1; Low: rate difference ≤ 0.1 and copy number difference ≤ 1. **(D)** Box plots showing the copy number variation of genes involved in the degradation of fucosylated and sialylated HMOs in persistent versus non-persistent strains across Westernized and non-Westernized infant populations, (Kruskal-Wallis test, *p*<0.001: ***, *p*<0.01: **, *p*<0.05: *, *p*>0.05: ns); Associations between the persistence rate of metagenome-assembled genomes (MAGs) in infants from **(E)** Westernized and **(F)** non-Westernized populations and the copy number of genes involved in the degradation of 2-fucosylated (GH29), 3-fucosylated (GH95), and sialylated (GH33) HMO; **(G)** Heatmaps of CAZy genes related to human milk oligosaccharide (HMO) degradation in *Bacteroides*, *Phocaeicola*, and *Parabacteroides*.

### Western and Non-Western Infants Harbor Distinct Cohabiting Microbiota for *B. longum* Subspecies

The presence of *Bl. longum* was negatively correlated with the presence of *Bl. infantis* in infants from Bangladesh, Sweden, Singapore, South Africa, Mozambique, and Zimbabwe (Fig. 5A). Additionally, a negative correlation was observed between *Bl. longum* and *Bl. iuvenis/suis* in infants from South Africa, Kenya and Zimbabwe. The presence of *Bl. infantis* in the infant was positively correlated with the presence of *Bl. iuvenis/suis* in populations from Bangladesh, Mozambique, Zimbabwe, and the Hadza community—groups associated with Westernized lifestyles (Fig. 5B). In Denmark and USA, *Bl. longum* was negatively correlated with *Bl. novel*. In addition, *Bl. longum* showed positive correlations with most species from the genera *Bacteroides*, *Phocaeicola*, *Parabacteroides*, and *Ruminococcus_B*, whereas *Bl. infantis* predominantly correlated negatively with most species within these same genera (Fig. 5B). The presence of *Bl. iuvenis/suis* was positively correlated with the presence of *Bl. infantis* in infants from Bangladesh, Mozambique, Kenya, and the Hadza community, while was negatively correlated with the presence of *Bl. infantis* in infants from Zimbabwe, Kenya, and South Africa (Fig. 5C).

**Fig. 5.**
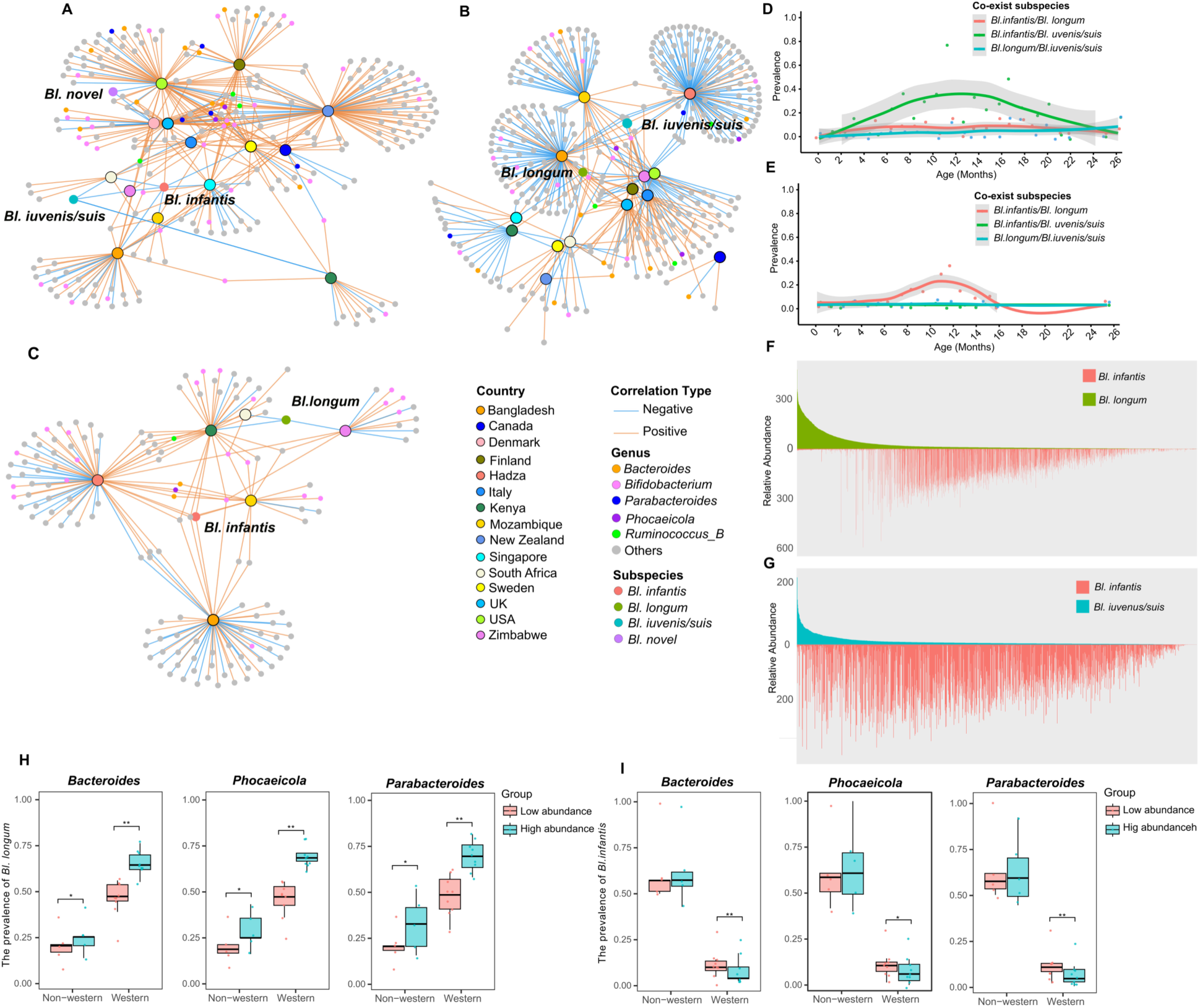
Correlation of *B. longum* Subspecies with Co-occurring HMO-Utilizing Bacteria in Early-Life Gut Microbiota with Western and Non-Western Lifestyles. Network of species significantly correlated with *Bl. longum* (A), *Bl. infantis* (B), and *Bl. iuvenis/suis* (C) across geographic cohorts. Country-labeled nodes (e.g., Denmark) denote the corresponding *Bl. longum*, *Bl. infantis*, or *Bl. iuvenis/suis* populations in infant cohorts from each geographic location; The prevalence of individuals harboring different combinations of two *B. longum* subspecies in infants with **(D)** non-western and **(E)** western lifestyles; (F) *Bl. infantis* vs. subsp. *longum* co-abundance within individual samples; (G) *Bl. infantis* vs. subsp. *iuvenis/suis* co-abundance within individual samples; Detection rates of (H) *Bl. longum* and (I) *Bl. infantis* in relation to high and low abundance states across *Bacteroides*, *Phocaeicola*, and *Parabacteroides*. Samples with an abundance above the overall median were defined as high-abundance samples, while those below the median were defined as low-abundance samples. * *p* < 0.05; ** *p* < 0.01; *** *p* < 0.001 by Mann-Whitney U test.

In non-Western infants, the co-occurrence prevalence of *Bl. infantis* and *Bl. iuvenis/susi* rose after birth, peaking at ∼12 months (60∼80% prevalence rate; Fig. 5D). Rates were significantly higher in infants aged 3–12 months compared to those <2 months or >12 months (Fig. S5A, Kruskal-Wallis test, *p* < 0.001). In contrast, co-occurrence between *Bl. infantis* and *Bl. longum*, or between *Bl. longum* and *Bl. iuvenis/suis*, was consistently very low (Fig. 5D; Fig. S5A). In Western infants, co-occurrence between *Bl. infantis* and *Bl. iuvenis/suis*, or between *Bl. longum* and *Bl. iuvenis/suis* was rarely observed throughout the first two postnatal years (Fig. 5E; Fig. S5B). For *Bl. infantis* and *Bl. longum*, co-occurrence was observed in only a small subset of infants at 8–16 months of age, though prevalence remained low (10∼20% prevalence rate; Fig. 5E; Fig. S5B). In addition, co-occurrence between *Bl. novel* and other subspecies are rare. Given the negative correlation between *Bl. longum* and *Bl. infantis* identified by co-occurrence network analysis, we further investigated the correlation patterns of abundances among *B. longum* subspecies. Our analysis of inter-subspecies abundance correlations revealed a striking inverse relationship between *Bl. longum* and *Bl. infantis* in the infant gut (Fig. 5F), whereas no significant negative correlation was observed between *Bl. iuvenis/suis* and *Bl. infantis* (Fig. 5G). In Western infants, *Bl. infantis* abundance was significantly higher in *Bl. longum*-negative than in *Bl. longum*-colonized individuals (Mann-Whitney U test, *p* < 0.05, except Mozambique cohort; Fig. S5C). In contrast, *Bl. infantis* abundance did not differ significantly between *Bl. iuvenis/suis*-negative and *Bl. iuvenis/suis*-colonized individuals (Fig. S5D). In Western infants, the abundances of *Bacteroides* (*p* < 0.05, except Canada, Italy, Singapore), *Phocaeicola*, and *Parabacteroides* (Fig. S5E) were significantly lower in *Bl. longum*-negative than in *Bl. longum*-colonized individuals. Conversely, *Bl. infantis*-negative infants showed significantly higher abundances of *Bacteroides* (*p* < 0.05, UK), *Phocaeicola* (*p* < 0.05, Italy and Singapore), and *Parabacteroides* (*p* < 0.05, Finland) compared to *Bl. infantis*-colonized infants (Fig. S5F). The high abundance infants (relative abundance above the national median) for *Bacteroides*, *Phocaeicola*, and *Parabacteroides* showed a significantly higher (*p* < 0.05) prevalence of *Bl. longum* (Fig. 5F). Conversely, this same high-abundance group exhibited a significantly lower (*p* < 0.05 in Western lifestyle country) prevalence of *Bl. infantis* (Fig. 5G). Further analysis revealed that, irrespective of infant lifestyle, the number of bacterial species within the genera *Bacteroides*, *Phocaeicola*, and *Parabacteroides* was significantly better in individuals harboring *Bl. longum* than in those lacking it (Mann-Whitney U test, *p* < 0.05, except Singapore, Kenya, and Mozambique cohorts, Fig. S5G, H).

### Distinct HMO utilization strategies drive the differential dominance of *Bl. longum* and *Bl. infantis* in infants across lifestyles

Analysis of the primary HMO-metabolizing bacteria in the gut of the infant revealed that species from genus *Bifidobacterium* mainly employ an intracellular degradation strategy (Fig. 6A). Conversely, those belonging to genera associated with western lifestyle, *Bacteroides*, *Phocaeicola*, and *Parabacteroides,* utilize an extracellular approach. Notably, *Ruminococcus_B gnavus* exhibits the capacity for both extracellular and intracellular HMO metabolism. An examination of HMO-metabolizing genes (fucosylated HMO, sialylated HMO) absent in *Bl. longum* revealed a clear phylogenetic dichotomy between the HMO-utilizing bacteria of infants from Western and non-Western lifestyles. With the exception of a few genes encoding extracellular enzymes in *B. bifidum*, key glycoside hydrolases (GH29, GH95, GH33, GH151) for fucosylated and sialylated HMOs were functionally intracellular in other *Bifidobacteria*, including *Bl. infantis*, *B. breve*, *B. kashiwanohense*, *B. pseudocatenulatum*, and *B. dentium*. In stark contrast, the homologous genes in *Bacteroides*, *Phocaeicola*, and *Parabacteroides* species were almost exclusively predicted to encode extracellular enzymes (Fig. 6A). Further quantification of microbial genes for fucosylated HMO and sialylated HMO metabolism highlighted a key dichotomy: a greater number of Western infants exhibited extracellular association of GH29, GH95, and GH33 genes, while a greater number of non-western infants exhibited intracellular association of these genes (Fig. 6B).

**Fig. 6.**
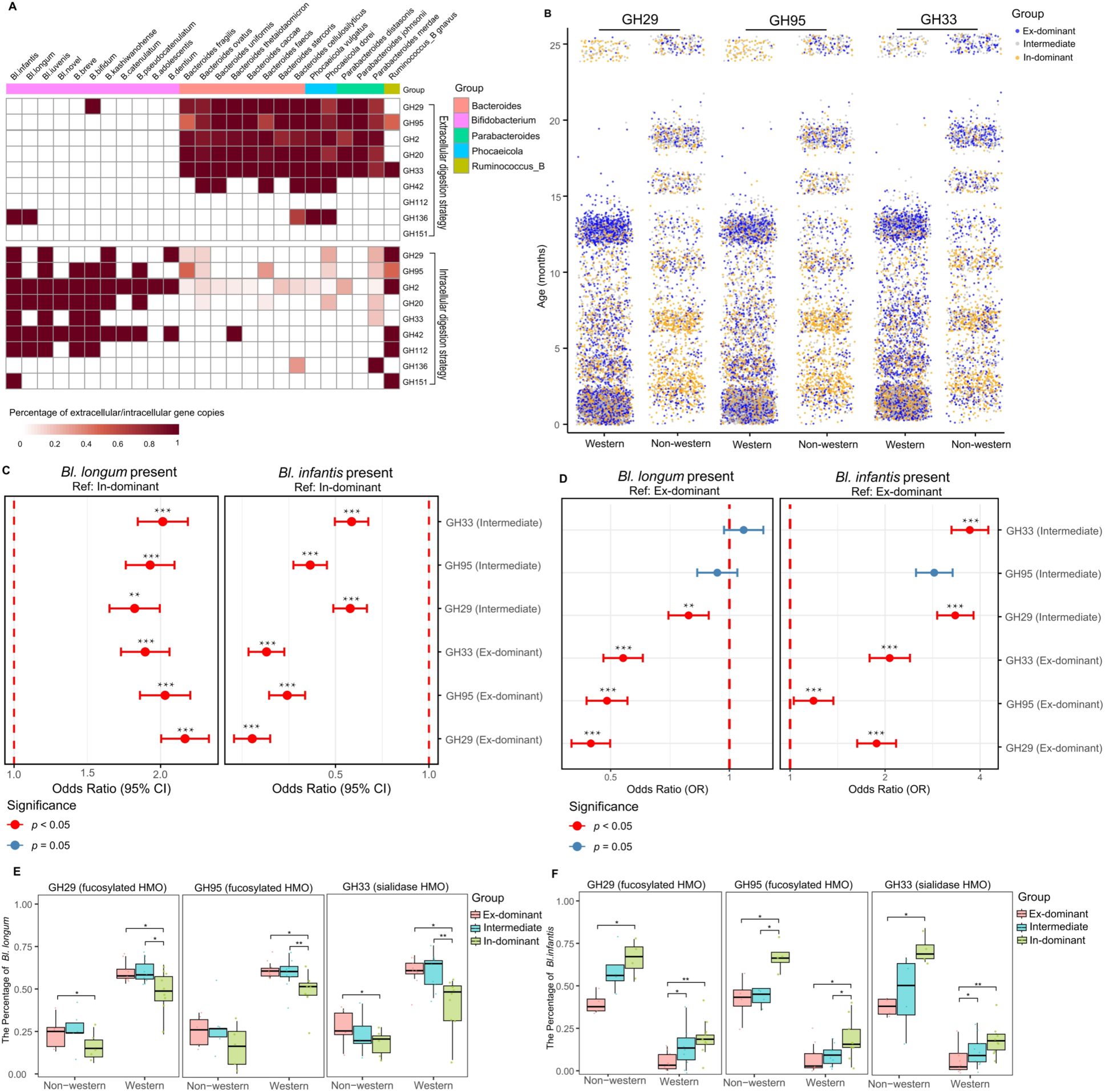
Comparative Analysis of Human Milk Oligosaccharide (HMO) Degradation Strategies in infants from Western and Non-Western Lifestyles. **(A)** Percentage of extracellular vs. intracellular HMO-metabolizing genes across key genera, values represent the proportion of identified genes per genus assigned to each localization category; **(B)** Samples were classified into three groups (Ex-dominant, In-dominant, and Intermediate) according to the relative abundance of extracellular vs. intracellular GH29, GH95, and GH33 genes in non-western and western lifestyles infants; Results from logistic regression models with In-dominant **(C)** or Ex-dominant **(D)** samples set as the reference level. Detection rates of (E) *Bl. longum* and (F) *Bl. infantis* in samples stratified by gene (GH29, GH95, GH33) and further categorized into Ex-dominant, In-dominant, or Intermediate groups based on extracellular vs. intracellular gene abundance. Ex-dominant: log₂ (Extracellular abundance/Intracellular abundance) ≥ 1; In-dominant: log₂ (Extracellular abundance/Intracellular abundance) ≤ -1; Intermediate: -1 < log₂ (Extracellular abundance / Intracellular abundance) < 1. * *p* < 0.05; ** *p* < 0.01; *** *p* < 0.001.

When In-dominant samples were used as the reference (Fig. 6C), Ex-dominant and intermediate samples showed opposite associations with *Bl. infantis* and *Bl. longum*: they were positively associated with the presence of *Bl. longum* (all OR > 1, *p* < 0.001, logistic regression models) but negatively associated with the presence of *Bl. infantis* (all OR < 1, *p* < 0.001, logistic regression models). Conversely, using Ex-dominant samples as the reference (Fig. 6D), In-dominant samples exhibited inverse patterns, being negatively associated with *Bl. longum* (all OR < 1, *p* < 0.001, logistic regression models) and positively associated with *Bl. infantis* (all OR > 1, *p* < 0.001, logistic regression models). Notably, the detection rate of *Bl. longum* was significantly higher (Kruskal-Wallis test, *p* < 0.05) in infants with Ex-dominant GH29, GH95, and GH33 genotypes than in those with In-dominant genotypes among Western-lifestyle countries (Fig. 6E, Figs. S6A-C ). In non-Western-lifestyle countries, this difference remained significant for GH29 and GH33. Conversely, the detection rate of *Bl. infantis* was significantly higher (Kruskal-Wallis test, *p* < 0.05) in infants with In-dominant genotypes across all three genes compared to those with Ex-dominant genotypes, in both Western and non-Western countries (Fig. 6F, Figs. S6D-F).

### Westernized Microbiota were the Key Factor that facilitated the colonization of *Bl. Longum*

In the COPSAC_2010_ cohort, multivariable logistic regression analysis revealed that, among the factors examined (including antibiotic exposure during the first month of life, urban versus rural living environment, delivery mode, breastfeeding), the presence of *Bacteroides* and *Phocaeicola* showed the strongest positive association with the colonization of *Bl. longum* from birth to one month of age. Furthermore, only these microbial factors and the presence of siblings were significantly associated with *Bl. longum* colonization (Table 1). In the Bangladesh cohort at age two months, the presence of *Ruminococcus_B* was positively associated with the colonization of *Bl. longum*, whereas the presence of *Bl. infantis* showed a negative association (Table 2). Similarly to the COPSAC_2010_ cohort, neither breastfeeding pattern (EB/PB) nor antibiotic exposure were significantly associated with the early-life colonization of *Bl. longum* in this population.

**Table 1.** Multivariable Logistic Regression Analysis of Factors Associated with *Bl. longum* Colonization in the COPSAC 2010 Cohort.

| Factors | Odds Ratio | 95% CI | p value |  |
| --- | --- | --- | --- | --- |
| Antibiotic exposure for baby after birth (Yes/No) | 1.179 | (0.579, 2.453) | 0.652 |  |
| *Bacteroides* (Present/Absent) | 1.959 | (1.272, 3.02) | 0.002 | ** |
| *Phocaeicola* (Present/Absent) | 1.746 | (1.141, 2.672) | 0.010 | * |
| *Parabacteroides* (Present/Absent) | 1.555 | (0.958, 2.539) | 0.075 |  |
| *Ruminococcus_B* (Present/Absent) | 1.141 | (0.688, 1.914) | 0.613 |  |
| Living environment (Urban/Rural) | 0.977 | (0.677, 1.412) | 0.903 |  |
| Family Income level (Medium/Low) | 0.974 | (0.524, 1.799) | 0.934 |  |
| Family Income level (High/Low) | 0.775 | (0.405, 1.470) | 0.436 |  |
| Intrapartum antibiotic exposure for mother during birth (Yes/No) | 0.648 | (0.361, 1.154) | 0.141 |  |
| Delivery (C-Section/Vaginal) | 1.637 | (0.854, 3.173) | 0.140 |  |
| Race (Caucasian/Non-caucasian) | 0.666 | (0.292, 1.494) | 0.325 |  |
| Sibling (Yes/No) | 1.545 | (1.068, 2.239) | 0.021 | * |
| Smoking during pregnancy (Yes/No) | 0.817 | (0.407, 1.645) | 0.567 |  |
| Breast (EB/NB) | 0.691 | (0.335, 1.379) | 0.085 |  |
| Breast (PB/NB) | 0.455 | (0.183, 1.104) | 0.528 |  |
| Ages | 0.990 | (0.961, 1.02) | 0.528 |  |

EB: Exclusive breast feeding; PB: Partial breastfeeding; NB: No breast feeding; OR >1: indicates a positive effect (increased odds of *Bl. longum* presence); OR<1: indicates a negative effect (increased odds of *Bl. longum* presence); Age (sample date minus birth date) was entered as a continuous covariate in the adjusted model. \* <0.05, \*\* < 0.01.

**Table 2.** Multivariable Logistic Regression Analysis *longum* Colonization in the Bangladesh Infant Cohort.

| Factors | Odds Ratio | 95% CI | p value |  |
| --- | --- | --- | --- | --- |
| *g_Bacteroides* | 0.93 | (0.18, 3.59) | 0.924 |  |
| *g_Phocaeicola* | 0.6 | (0.04, 4.76) | 0.668 |  |
| *g_Parabacteroides* | 3.37 | (0.39, 24.63) | 0.234 |  |
| *g_Ruminococcus_B* | 6.94 | (1.40, 40.18) | 0.0195 | * |
| *Bl. infantis* | 0.35 | (0.16, 0.76) | 0.0076 | ** |
| Delivery (C-Section/Vaginal) | 1.73 | (0.66, 5.40) | 0.2966 |  |
| Breast (EB/NB) | 0.00 | NE | 0.9951 |  |
| Breast (PB/NB) | 1.39 | (0.07, 10.15) | 0.7779 |  |
| Antibiotic exposure for baby after birth<br>(Yes/No) | 0.00 | NE | 0.9903 |  |

EB: Exclusive breast feeding; PB: Partial breastfeeding; NB: No breast feeding; OR >1: indicates a positive effect (increased odds of *Bl. longum* presence); OR<1: indicates a negative effect (increased odds of *Bl. longum* presence); NE: Not estimable due to perfect separation or sparse data. \* <0.05, \*\* < 0.01.

### *Bl. longum*, but Not *Bl. infantis*, Collaborates with Westernized Microbiota *in vitro*

To confirm the observational findings of HMO utilization, an *in vitro* experiment was conducted to assess the growth of the *B. longum* subspecies in 2′-FL (Fig. 7A). Consistent with genomic predictions indicating a limited repertoire of fucosylated HMO utilization genes, *Bl*. *longum* exhibited negligible growth when cultured alone on 2′-FL as the sole carbon source, whereas *Bl*. *infantis* and *Bacteroides fragilis* grew well under identical conditions (Fig. 7B). In pairwise co-culture experiments, *Bl*. *infantis* failed to promote the growth of *Bl*. *longum* on 2′-FL (Fig. 7C), while *B. fragilis* enhanced *Bl*. *longum* growth (Fig. 7D). During the first 36 hours of co-culture with *B. fragilis*, *Bl*. *infantis* displayed slower growth compared to monoculture (Fig. 7E). Sequential inoculation experiments further demonstrated that pre-culturing *Bl*. *infantis* for 12 hours did not enhance *Bl*. *longum* growth (Fig. 7F), whereas pre-culturing *B. fragilis* significantly promoted it (Fig. 7G). Compared to monocultures of *Bl*. *infantis* and *B. fragilis*, sequential cultivation where *Bl*. *infantis* was cultured first followed by the introduction of *B. fragilis* resulted in the inhibition of growth for both (Fig 7H). Conversely, when *B. fragilis* was cultured first, the growth of *Bl*. *infantis* was also inferior to that in its monoculture (Fig 7I). Similarly, when conditioned media derived from 24-hour monocultures were filtered and inoculated with *Bl*. *longum*, growth curves (Fig. 7J) and absolute abundances (Fig. 6K) were markedly higher in *B. fragilis*-conditioned medium compared to *Bl*. *infantis*-conditioned medium. In contrast, *B. fragilis*-conditioned medium slightly supported the growth of *Bl. infantis*, whereas *Bl*. *infantis*-conditioned medium did not support *B. fragilis* (Fig. 7L, M). Together, these results demonstrate that *B. fragilis*, but not *Bl*. *infantis*, facilitates *Bl*. *longum* growth on 2′-FL under *in vitro* conditions.

**Fig. 7.**
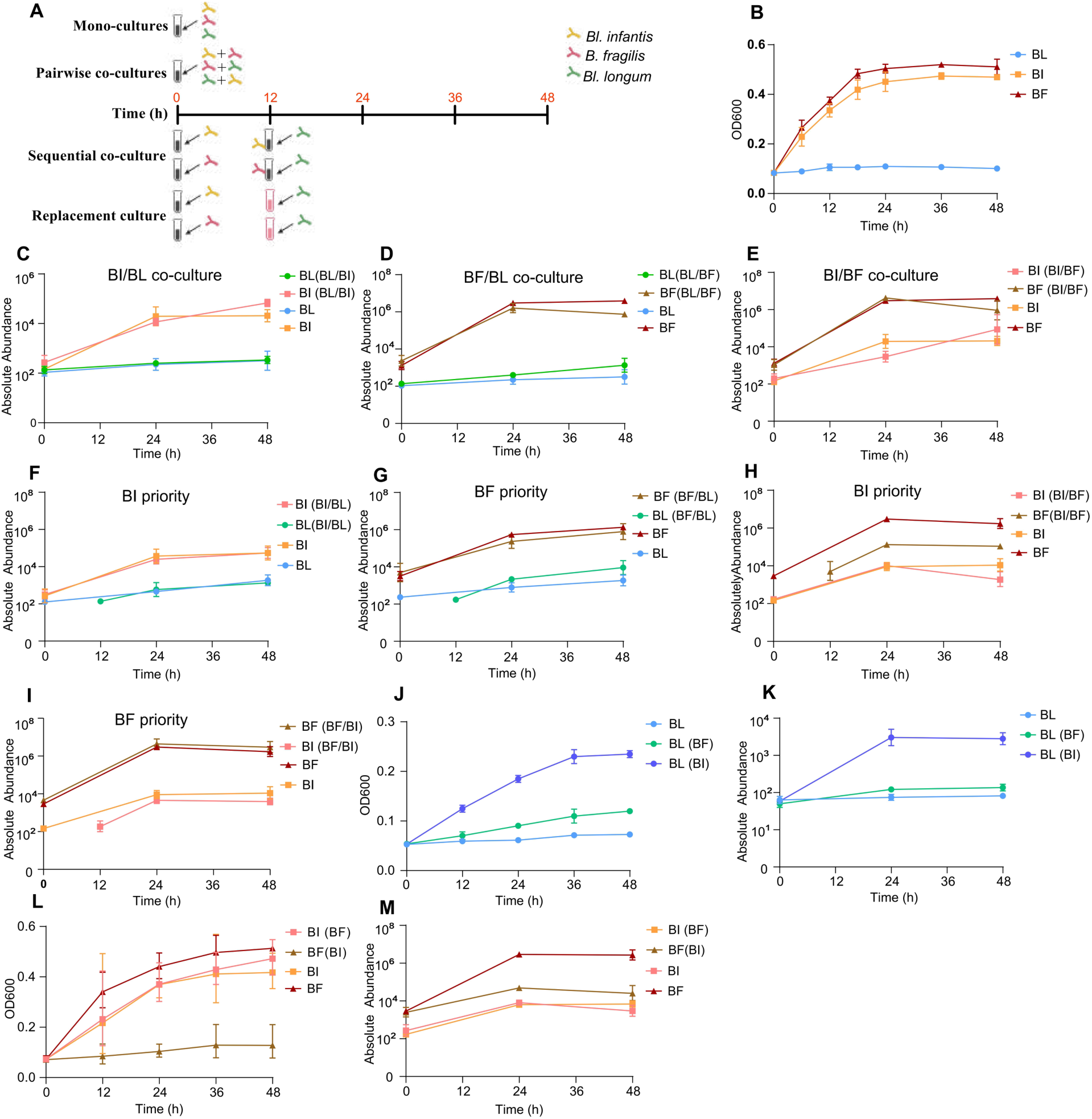
Pairwise cultivation on 2-FL as the sole carbon source. **(A)** Experimental design; **(B)** Monoculture growth curves of each strain in 2′-FL-supplemented PYG medium, monitored by OD600 readings over time; **(C-E)** Culturing results of strain pairs *Bl. longum*/*Bl. infantis*, *Bl. longum*/*B. fragilis*, and *Bl. infantis*/*B. fragilis* in monoculture vs co-culture; **(F)** Culturing results of *Bl. longum* and *Bl. infantis* in monoculture versus sequential co-culture (*Bl. infantis* pre-cultured for 12 h); **(G)** Culturing results of *Bl. longum* and *B. fragilis* in monoculture versus sequential co-culture (*B. fragilis* pre-cultured for 12 h); **(H)** Culturing results of *Bl. infantis* and *B. fragilis* in monoculture versus sequential co-culture (*Bl. infantis* pre-cultured for 12 h); **(I)** Culturing results of *Bl. infantis* and *B. fragilis* in monoculture versus sequential co-culture (*B. fragilis* pre-cultured for 12 h); **(J)** Growth curves and **(K)** absolute abundance of *Bl. longum* in *Bl. infantis*-conditioned or *B. fragilis*-conditioned medium versus monoculture; **(L)** Growth curves and **(M)** absolute abundance of *Bl. infantis* (or *B. fragilis*) in *B. fragilis* (or *Bl. infantis*)-conditioned medium versus monoculture; *Bl. infantis*-conditioned: *Bl. infantis* pre-cultured for 24 h, then removed by filtration; *B. fragilis*-conditioned: *B. fragilis* pre-cultured for 24 h, then removed by filtration. BI: *Bl. infantis*; BL: *Bl. longum*; BF: *B. fragilis*;

## Discussion

Our study expands the known diversity of *B. longum* in the infant gut by revealing the presence of five distinct subspecies. This finding markedly expands our understanding of subspecies-level diversity within this species in early life, as previous studies have consistently reported only three or four subspecies in the same niche ^3,8^. Although *Bl. iuvenis* and *Bl. suis* exhibit high genomic similarity ^9^, they remain distinct subspecies, meeting established delineation criteria based on ANI-defined genomic divergence (≥2%) and differences in metabolic gene profiles. Previous studies have generally classified these two taxa as a single subspecies ^3,8,15^, likely due to the limited inclusion of diverse cohorts, insufficient recovery of metagenome-assembled genomes (MAGs), and the lack of comparative analyses incorporating non-human-derived *Bl. suis*. A recent genomic study has proposed *Bl. infantis* as an independent species ^8^; however, its ANI divergence exceeds 5% only with *Bl. longum* and *Bl. novel*, while its divergence from *Bl. iuvenis* is even smaller than that between *Bl. novel* and *Bl. iuvenis*. Therefore, its taxonomic status as a separate species remains unresolved.

Notably, the close phylogenetic relationship between ancient Mexican *Bl. longum* and contemporary East Asian lineages are consistent with the widely accepted model of Asian ancestry in Indigenous American populations ^16–18^. The estimated divergence time between modern Asian and ancient American *B. longum* strains (∼11,743 years; 95% HPD: ∼2,400–30,400 years) suggests a long-term association between this gut symbiont and human populations. Notably, the upper bound of the 95% HPD interval overlaps with the proposed timeframe for the initial peopling of the Americas via the Bering Land Bridge. Current archaeological and palaeogenomic evidence suggests that the ancestors of Native Americans diverged from Siberian populations sometime between ∼23,000 and ∼15,000 years ago ^18–20^, with some archaeological sites in Mexico (e.g., Chiquihuite Cave) yielding evidence of human occupation as early as ∼33,000 years ago ^21^. This temporal concordance is consistent with a shared demographic history between humans and *B. longum*, supporting the use of ancient gut commensals as complementary “molecular fossils” for reconstructing human migration history.

While previous extensive studies have not detected the prevalence of *B. longum* in non-human primates ^22–24^. Here, our multi-host comparative analyses not only confirm this absence but further reveal that *B. longum* is prevalent in livestock, particularly bovine, and only detectable in more recent ancient human samples. It indicates that the establishment of *B. longum* as a common member of the human gut microbiome is a relatively recent event in our evolutionary history. In addition, human-derived MAGs (genome) exhibited closer phylogenetic relationships with those from domesticated animals in the same geographic region, suggesting that *B. longum* is probably not an ancestral primate symbiont but was likely acquired through cross-species transmission associated with animal domestication. However, human-derived *Bl. suis* utilizes HMOs more effectively than animal-derived strains, suggesting that *B. longum* may have evolved an HMO-preferring phenotype after transfer from animals to humans. This evolutionary perspective provides important context for the present-day distribution of *B. longum* subspecies, indicating that their ecological roles may reflect not only contemporary lifestyle factors but also relatively recent host–microbe associations events.

The introduction of bacterial communities from domesticated animals to humans is not an isolated event ^25^. Previous studies have confirmed that the genus *Ruminococcus*, which possesses cellulose-metabolizing functions in the human gut, may have originated from ruminants ^26^. However, the widespread prevalence of this genus in non-human primates makes it difficult to determine whether *Ruminococcus* originated via vertical transmission from human ancestors or cross-species transmission from ruminants. Furthermore, previous studies have mainly explored this evolutionary relationship at the species level ^27^. Given that different subspecies within the same species often exhibit host specificity ^28,29^, investigating species origins at the subspecies level is more scientifically meaningful. Our study provides subspecies-level evidence suggesting that some *B. longum* subspecies may have arisen through a host transition between humans and domestic animals.

A central insight emerging from the data is the asymmetric response of *B. longum* subspecies to lifestyle-associated microbiota configurations. While *Bl. longum* and *Bl. novel* exhibits minimal phylogenetic structuring by lifestyle, the HMO-specialized clades (*Bl*. *infantis*, *Bl*. *iuvenis*, and *Bl*. *suis*) show strong lifestyle-linked divergence. Among *Bl. infantis* strains, three major phylogenetic clades were identified, corresponding to a Western lifestyle-associated Chinese clade, a Southern African clade, and a South Asian/East African clade, consistent with the population structure previously reported by Shao and colleagues ^8^. Notably, strains originating from China consistently clustered with those from Westernized infant populations, suggesting that contemporary Chinese lifestyles may be undergoing progressive Westernization at the microbiome level ^30^. This pattern argues against a purely dispersal-driven explanation and instead suggests that selective pressures act differentially on subspecies whose fitness may depend on access to specific carbohydrate niches. The absence of pronounced metabolic divergence within *Bl. longum* across lifestyles further supports this interpretation. This may be explained by *Bl. longum* having a wider host age range and greater niche breadth ^31^, unlike *B. infantis*, *Bl*. *iuvenis*, and *Bl*. *suis*, which is mainly prevalent during infancy. This enables the *Bl. longum* to develop a broader genotype for environmental robustness. In contrast, different lifestyles may more readily shape adaptive evolution in those infants-specialized subspecies, given its experience of more selective pressures during early life.

By resolving the distribution of *B. longum* subspecies across diverse infant populations, this study highlights how early-life colonization is shaped by interactions between microbial metabolic strategies and community context, rather than solely by host lifestyle. The results go beyond the long-standing assumption ^32,33^ that breastfeeding, delivery mode, and exposure to antibiotics are the determinants for the success of HMO-specialist *B. longum*, and instead point to community-mediated selection as a dominant force structuring the persistence and prevalence of *B. longum* subspecies. These differences are particularly salient when viewed alongside age-dependent dynamics, suggesting that subspecies are differentially tuned to specific windows of infant dietary ecology. Importantly, these distinctions persist across cohorts, indicating that subspecies identity captures a stable ecological strategy that interacts with, rather than passively reflects, host and environmental variation. Therefore, this study helps explain the lifestyle-stratified dominance of *B. longum* subspecies. For instance, *Bl*. *infantis* is depleted in infants with a Western lifestyle ^34^. The functional comparisons reinforce that *B. longum* subspecies represent distinct ecological strategies rather than stochastic variations. A major contribution of this work lies in linking *B. longum* subspecies success in distinct lifestyle to the metabolic profiles of cohabiting microbial communities. The consistent association of *Bl*. *longum* with taxa such as *Bacteroides*, *Phocaeicola*, and *Parabacteroides*, alongside its negative association with *Bl. infantis* across multiple populations suggests that cross-feeding, competition or niche displacement occurs at the level of catabolic mechanisms rather than substrate availability. These associations were consistent across cohorts and are supported by regression analyses that identify microbiota features, rather than feeding mode or antibiotic exposure, as the strongest determinants of early-life *Bl. longum* colonization. The ability of *Bl. longum* to engage in cross-feeding interactions with other gut community members for carbohydrate utilization ^35^ may represent a key determinant of its capacity to adapt to a broader host age range and more diverse geographical populations.

The analysis of HMO-degrading enzyme localization offers a mechanistic explanation for these patterns. In *Bifidobacteria*, the dominant genus in infants from non-westernized lifestyles ^2^, key enzymes involved in fucosylated and sialylated HMO metabolism are predominantly intracellular, consistent with a “private goods” strategy ^36^ in which intact HMOs are imported and metabolized within the cell. In contrast, homologous enzymes in dominant Western-associated taxa are mostly extracellular ^37^, enabling cleavage of HMOs in the lumen and release of shared breakdown products. This distinction has important ecological consequences: extracellular degradation can decouple HMO availability from competitive advantage for HMO specialists, broadening access to derived sugars and reshaping selective pressures. Within this framework, the success of *Bl*. *longum* in Westernized microbiotas may reflect its compatibility with an ecosystem dominated by extracellular HMO processing and cross-feeding networks, rather than superior intrinsic HMO metabolism. Conversely, *Bl*. *infantis*, which relies on efficient intracellular uptake of intact HMOs, may lose its selective advantage when HMOs are extensively processed outside the cell. In this study, in vitro pairwise co-culture experiments on 2-FL as the sole carbon source further confirmed that the distinct HMO utilization patterns of *Bl. infantis* and *Bl. longum* underlie their differential prevalence in non-Western versus Western lifestyles. This interpretation provides a coherent explanation for the repeated observation that exclusive breastfeeding does not reliably restore *Bl*. *infantis* in Western cohorts, despite abundant HMO supply.

The *Bl.* “*novel*” clade ^5^ further underscores how lifestyle-associated niches may select for lineages with distinct metabolic priorities. Its limited HMO repertoire, enrichment in starch-related genes, and association with early infancy and cesarean delivery suggest adaptation to alternative carbohydrate landscapes and transmission routes common in Western settings. Whether this clade represents a transient colonizer, a reservoir for later adaptation, or a lineage favored under specific dietary or medical exposures remains an open question that warrants targeted isolation and phenotypic characterization.

These findings have direct implications for microbiome-based interventions. Strategies aimed at restoring *Bl. infantis* in Western populations have largely focused on strain supplementation or increased HMO provision. The present data suggest that such approaches may fail if they do not account for community-level metabolic context. Restoring HMO specialists may require reshaping substrate partitioning and microbial interactions, for example by promoting bifidobacterial consortia that reinforce intracellular HMO utilization or by limiting competitive extracellular degradation during critical colonization windows.

## Method Details Ethics

This COPSAC_2010_ study was approved by the Ethics Committee of the Capital Region of Denmark (ref. H-B-2008-093) and the Danish Data Protection Agency (ref. 2015-41-3696), and conducted in accordance with the Declaration of Helsinki. Written and verbal informed consent were obtained from all parents prior to participation.

### Experimental Model and Subject Details

In this study, we included fecal samples from participants of the COPSAC_2010_ cohort. The COPSAC_2010_ mother-child cohort is a population-based longitudinal clinical study involving 736 pregnant women and their 700 children. Fecal samples used in the present study were collected at the ages of one month, and one year. Fecal samples were collected in sterile plastic containers, transported to Statens Serum Institut (Copenhagen, Denmark), and upon arrival, mixed with 1 mL of 10% (vol/vol) glycerol broth (SSI, Copenhagen, Denmark). The mixtures were then frozen at – 80 °C until further use. Additionally, clinical measurements and environmental exposures metadata were recorded prospectively starting from the 24th week of gestation. Through this collaboration, two longitudinal infant cohorts were added: a Bangladesh cohort of 267 infants from healthy mothers, followed from birth to age two in a peri-urban Dhaka community ^3^, and a Swedish cohort of 98 mother-child pairs with fecal samples collected at one week, four months, and one year ^38^.

### DNA Extraction and Shotgun Metagenomic Sequencing of 1-Month Fecal Samples

Previous studies have conducted metagenomic sequencing on stool samples collected from 1-year-old infants within the COPSAC_2010_ cohort ^39^. Total DNA was extracted from 200 mg of 1-month fecal material using the Kapa Hyper Prep kit (KAPA Biosystems, Wilmington, MA, USA) according to the manufacturer’s instructions. The concentration and purity of DNA were measured using a Qubit fluorometer (Thermo Fisher Scientific, Waltham, MA, USA), and agarose gel electrophoresis was conducted to evaluate DNA quality. Only samples meeting the standards for metagenomic sequencing proceeded to library preparation. DNA libraries were prepared for metagenomic sequencing on the Illumina Novaseq platform with the TruSeq DNA, producing 150 bp paired-end reads.

### Extended Cohorts and Lifestyle Classification

To assess the geographic distribution and genetic variation of *B. longum* subspecies, metagenomic sequencing reads from infant stool samples (ages 0–2 years) across 18 distinct studies conducted in various regions (n = 6,919) were retrieved from publicly available repositories (Table S7). Samples were categorized by lifestyle based on classifications from the U.N. Human Development Index (HDI, https://hdr.undp.org/content/human-development-report-2021-2022). Industrialized infants are typically from urban areas in countries with very high or high human development (HDI ≥ 0.9), as classified by the HDI. These infants are generally exposed to high-calorie diets, frequent consumption of formula milk and processed foods, higher rates of cesarean births, increased exposure to antibiotics or antimicrobials, and limited breastfeeding and animal contact. These factors reflect the infrastructure, healthcare, and educational access typical of industrialized societies. In contrast, non-industrialized infants are from underdeveloped regions classified as medium or low human development (HDI < 0.7). They are more likely to experience extended breastfeeding, greater exposure to animals, reliance on locally produced foods, and limited access to pharmaceuticals, shaping a lifestyle influenced by local resources and traditional practices.

According to these definitions, in this study, Bangladesh ^3^ Zimbabwe ^40^, Kenya ^13^ South Africa ^41^, Mozambique ^42^ Ethiopia ^43^, and Tanzania ^2^ were classified as non-industrialized populations, with HDI scores ranging from 0.461 to 0.67. In contrast, Denmark, Sweden, Finland ^44^ the United Kingdom ^45^ New Zealand ^46^ Singapore ^47^ Italy ^48,49^, Canada ^50^, and the United States^42,51,52^ was classified as industrialized populations, with HDI scores exceeding 0.9 (Table S7). Given that China’s Human Development Index (HDI) ranges between 0.7 and 0.9, the Chinese infant cohorts ^53^ were not classified into either Western or non-Western lifestyle groups.

### Cattle, Non-human Primate, and Ancient Human Data

Fecal metagenomic datasets from cattle were obtained from five independent studies (Table S8), while non-human primate fecal metagenomes were collected from 15 independent studies encompassing 37 species (Table S9). In addition, 72 ancient human fecal or gut metagenomes, spanning approximately 150 to 5,300 years before present, were also retrieved from publicly available repositories (Table S10).

### Metagenomics data processing

Adapter sequences were removed using BBDuk v39.01 with the following parameters: “ktrim=r k=23 mink=11 hdist=1 hdist2=0 ptpe tbo” (https://sourceforge.net/projects/bbmap/). Sequences with low quality (< 20) and those shorter than 100 bases are trimmed using Sickle v1.33 ^54^. To remove human host sequence contamination, all reads mapped to the human genome (hg19) were removed using BBMap v38.9 with default settings ^55^. Clean reads were assembled into contigs using SPAdes v3.15.5, with k-mer sizes set to 21, 33, 55, 77, 99, and 127 ^56^. Contigs longer than 1,500 bases were used for binning metagenome-assembled genomes (MAGs) for each sample using MetaWRAP v1.3.2 ^57^ and VAMB v3.0.8 ^58^. The binning module of MetaWRAP employed three metagenomic binning tools—MaxBin2, metaBAT2, and CONCOCT—to enhance the quality of the binning process. Genome completeness and contamination were evaluated using CheckM v1.2.2 ^59^. High-quality MAGs, defined as having at least 95% completeness and less than 5% contamination, were retained for further analysis. Taxonomic assignments for each MAG were performed using the GTDB-tk v2.1 according to Genome Taxonomy Database ^60^

### Evolutionary and Phylogenetic Insights into *B. longum*

A total of 1,591 publicly available *B. longum* genomes were retrieved from the National Center for Biotechnology Information (NCBI) (Table S11). dRep v3.4.3 ^61^ was used to dereplicate *B. longum* MAGs and publicly available *B. longum* genomes within each cohort or from NCBI, applying a similarity threshold of 99%. Phylogenetic analyses of the dereplicated *B. longum* publicly genomes and MAGs were performed with PhyloPhlAn v3.0.68 ^62^ The phylogeny tree was built using 400 universal marker genes identified by PhyloPhlAn, with the parameters set to: “--diversity low --fast”. The set of external tools, along with their specified options, was provided through the configuration file references_config.cfg.

- DIAMOND v2.0.13 ^63^ was used for nucleotide-based mapping with “Blastx” and amino acid-based mapping with “Blastp,” both executed with the parameters “--quiet --more-sensitive --id 50 --max-hsps 35 -k 0”.
- MAFFT v7.490 ^64^, with the parameters as “--quiet --anysymbol --auto”.
- trimAl v1.4.1 ^65^ with the parameters as “-gappyout”.
- FastTree v2.1.11 ^66^, with the parameters as “-quiet -pseudo -spr 4 -mlacc 2 -slownni -fastest - no2nd -mlnni 4 -lg”.
- RaxML v8.2.12 ^67^, with the parameters as “-p 1989 -m PROTCATLG”.

To elucidate the origin of human-associated *B. longum*, we constructed phylogenetic trees using the dereplicated *B. longum* MAGs from each country identified in this study, complete *B. longum* genomes from NCBI, as well as strains derived from animals and ancient humans. In parallel, we generated phylogenies for the human- and animal-associated *Bl*. *iuvenis* and *Bl. suis*. Both trees were rooted using *Bifidobacterium breve*.

To explore the evolutionary relationships of *B. longum* subspecies across different lifestyles, phylogenetic tree of core genes for each *B. longum* subspecies was also built with PhyloPhlAn based on core genes sets for each *B. longum* subspecies (100% present across all samples of a given subspecies) identified through Roary v3.13.0 ^68^ with a minimum gene identity threshold of 90%. The external tools were executed with the same parameters as previously described. Phylogenetic trees based on whole genomes were visualized using the ggtree package ^69^ in R v4.3.3, with *Bifidobacterium breve* set as the root. The phylogenetic tree of core genes for each B. *longum* subspecies were visualized using GraPhlAn v1.13 ^70^.

### Molecular Clock Analysis and Divergence Time Estimation

To estimate the divergence time between ancient American and modern Asian *Bl. longum* lineages, we performed Bayesian molecular clock analyses using BEAST v2.7.7 ^71^ based on a recombination-filtered core-genome alignment generated with Gubbins v3.4.3 ^72^. Sequence evolution was modeled under the HKY substitution model with four gamma-distributed rate categories. An uncorrelated lognormal relaxed molecular clock ^73^ was applied, with a mean substitution rate prior of 1.0 × 10⁻⁶ substitutions per site per year. A Yule speciation process was specified as the tree prior.

Temporal calibration was achieved by assigning radiocarbon ages of 1,200 and 1,400 years before present to the ancient Mexican genomes, whereas all contemporary genomes were assigned an age of zero. Markov chain Monte Carlo (MCMC) analyses were run for 50 million generations, sampling parameters every 10,000 generations. Convergence and mixing were assessed using Tracer v1.7.2 ^74^, and all parameters achieved effective sample size (ESS) values greater than 200.

The maximum clade credibility (MCC) tree was generated using TreeAnnotator v2.7.7 ^71^, from which the median divergence time and the corresponding 95% highest posterior density (HPD) interval for the most recent common ancestor (tMRCA) of the ancient American and modern Asian lineages were estimated.

### Genetic Distance between and within *B. longum* Subspecies

Pairwise average nucleotide identity (ANI) distances among and within the five *B. longum* clades were computed using pyani v0.2.11 with the ANIb method ^75^. Core genome alignments were generated via the integrated PRANK plugin in Roary v3.13.0 ^76^, followed by calculation of pairwise core genome distances from these alignments. Gene content similarities were assessed using the Roary-generated pangenome matrix, with pairwise Jaccard similarity coefficients quantified through the vegan R package.

### Construction of *B. longum* Marker Genes and Evaluation of *B. longum* Prevalence and Abundance

Open Reading Frames (ORFs) for the dereplicated *B. longum* public genomes and MAGs were predicted using Prodigal ^77^. The total *B. longum* pangenomes were determined using Roary with a 90% protein identity threshold ^68^. Markers genes of each subspecies of the *B. longum* were defined as present in >90% of a given subspecies but absent in all others. Reads from each metagenomic sample were mapped to subspecies-specific markers using Bowtie2 ^78^ with the default parameter. A marker gene was considered present if it had coverage ≥ 0.5X. A subspecies was considered present if ≥ 50% of its specific marker genes were detected. The relative abundance of *B. longum* subspecies was calculated using the following formula:

Abundance (subspecies) = (Mean subspecies marker genes coverage × Approximate genome size (bp)) / Total metagenome size (bp)

### Prevalence and Abundance of Non-*B. longum* MAGs

Non-*B. longum* MAGs from all cohorts were dereplicated using dRep v3.4.3 at a 98% similarity threshold ^61^. A non-redundant database of representative MAGs was generated following the inStrain tutorial ^79^, utilizing Bowtie2 and Prodigal. Reads from each individual fecal sample were mapped to the non-redundant database of representative MAGs using Bowtie2 as part of the inStrain (v1.3.4) *profile*. These mappings were processed with inStrain *profile* using default settings to generate strain profiles. MAGs with ≥ 0.5X coverage in samples were considered to be present. The relative abundance was calculated using the following formula:

Abundance (MAG) = (Average coverage depth of all scaffolds of this MAG) / Total metagenome size (bp).

### Construction of Non-Redundant Gene Sets for Each Cohort

Assembled contigs longer than 1,500 bp were retained for subsequent analyses. For each cohort, all the contigs were merged into a single file representing its specific cohort. Open reading frames (ORFs) were predicted using Prodigal ^77^. The non-redundant gene set of each cohort was generated with a 90% identity cutoff using CD-HIT (v4.8.1) ^80^. The reads were mapped against the merged contigs of each cohort using BWA (v0.1.79) ^81^. The resulting alignments were sorted and indexed with Samtools prior to coverage calculation. The coverage profiles were generated by running MetaBAT2’s depth summarization command “jgi_summarize_bam_contig_depths” ^82^. Contig-level relative abundance was computed as:

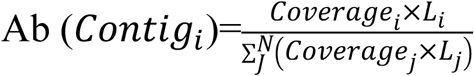

where Coverage_i_ is the mean depth for Contig *i* and *L*_i_ its length. To derive gene-level abundance, each gene’s proportional length within its parent contig was used to scale the contig relative abundance:

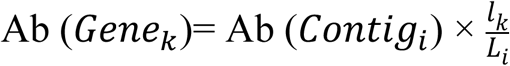

where *l*_k_ is the length of gene *k* on contig *i* (*L*_i_). This two-step normalization accounts for both sequencing depth and contig size, yielding accurate relative abundances for individual genes across samples.

### Carbohydrate active enzymes (CAZymes) annotation

CAZymes were found in the set of non-redundant public genomes and MAGs of *B. longum* using run_dbcan v4.1.4 with the dbCAN3 framework ^83^. Genomes (MAGs) were annotated by HMMER searches against the dbCAN CAZyme domain HMM database (v12) (E-value<1e-15, coverage>0.5) and DIAMOND was used to annotate query sequences with hits in the CAZy database (CAZyDB.07262023) (E-value<1e-102, coverage≥80%) ^84^. For discordant CAZyme annotations between HMMER and DIAMOND, HMMER predictions were prioritized. Non-*B. longum* MAGs and the non-redundant gene set of each cohort were annotated for CAZymes using identical methods and criteria.

### Genomic Map of Fucosylated HMO Transport and Metabolism Genes in the Five *B. longum* Subspecies

For each *B. longum* subspecies, the representative genomes with high completeness were selected for synteny analysis. Specifically, the genomes of *Bl. infantis* ATCC 15697, *Bl. iuvenis* NCC5000, human-derived *Bl. suis* YGMCC 0209, and animal-derived *Bl. suis* JCM 19995 was obtained from NCBI or the Integrated Microbial Genomes (IMG) system. Homologous biosynthetic gene clusters were identified using clinker (v0.0.31) ^85^. The five genomes were annotated using the RAST (Rapid Annotation using Subsystem Technology) server to map coding sequences (CDSs) to subsystems and reconstruct metabolic pathways ^86^. Subsequently, genes involved in human milk oligosaccharide (HMO) transport and metabolism were identified to compare homologous biosynthetic gene clusters across these genomes.

### Prediction of Extracellular or Intracellular Functions of HMO Metabolic Genes

Representative genomes of key HMO-metabolizing strains with high completeness were retrieved from NCBI and Integrated Microbial Genomes (IMG), including genera such as *Bifidobacterium*, *Bacteroidetes*, *Phocaeicola*, *Parabacteroides*, and *Ruminococcus_B* (Table S12). The annotation of CAZymes in the representative genomes was conducted as described above. Genes annotated as being related to HMO metabolism were subsequently predicted to function either extracellularly or intracellularly. Amino acid sequences were analyzed using SignalP 6.0 with default parameters^87^. The amino acid sequence was analyzed to identify the gene encoding the signal peptide. This gene was classified as encoding an extracellularly acting protein, whereas genes lacking a predicted signal peptide were considered to encode intracellular proteins. The signal peptides of HMO metabolic genes in all MAGs as well as the non-redundant gene set were predicted using the same method.

### Special Definition

Bacterial species detected within the same individual both before one month of age and at approximately 12 months of age are defined as persistent species, whereas those present before one month of age but absent at approximately 12 months of age in the same individual are classified as non-persistent species (ref Fig. 4A).

For each HMO utilization gene, differences between persistent and lost species were then computed as (ref Fig. 4B, C):

- Prevalence rate difference = prevalence rate (persistent) – prevalence rate (lost)
- Mean copy number difference = mean copy number (persistent) – mean copy number (lost)

Based on the joint distribution of these two-difference metrics, we classified HMO utilization gene into three levels to prioritize candidates for further interpretation:

- High: prevalence rate difference > 0.25 and mean copy number difference > 1;
- Medium: not meeting High criteria, but either prevalence rate difference > 0.1 or copy number difference > 1;
- Low: prevalence rate difference ≤ 0.1 and copy number difference ≤ 1. (i.e., meeting neither the high nor medium conditions).

Sample categorization (Ex-dominant, In-dominant, or Intermediate) was performed according to the relative abundance distribution of GH29, GH95, and GH33 genes in extracellular versus intracellular compartments. Specifically, we calculated the log₂ ratio of extracellular abundance to intracellular abundance for these genes. Samples were then assigned to one of three groups based on the following thresholds (ref Fig. 6C-F): (1) Ex-dominant: Samples with a log₂ ratio ≥ 1; (2) In-dominant: Samples with a log₂ ratio ≤ -1; (3) Intermediate: Samples with a log₂ ratio between - 1 and 1 (exclusive).

### In Vitro Bacterial Co-culture Experiments

To further clarify the interrelationship among *Bl*. *longum*, *Bl*. *infantis*, and the Westernized microbiota, we performed pairwise in vitro co-culture experiments involving *Bl*. *longum*, *Bl*. *infantis*, and *Bacteroides fragilis*. *Bl*. *infantis* (ATCC 15679), *Bl*. *longum* (ATCC 15707), and *B. fragilis* (ATCC 25285) were purchased from ATCC Company (Manassas, Virginia, United States). Prior to experiments, *B. fragilis* was expanded in Gifu Anaerobic Medium (GAM), while *Bl. longum* and *Bl. infantis* were cultured in Peptone-Yeast-Glucose (PYG) medium. All cultures were incubated under anoxic conditions using the Anaerobic Workstation (MiniMACS DG250; Don Whitley Scientific, UK) at 37 °C. Cultures were performed with five different patterns: mono-cultures, pairwise co-cultures, sequential co-culture, replacement culture. For sequential co-culture, *Bl*. *infantis* (or *B. fragilis*) was introduced into a 12-hour pre-established culture of *Bl*. *longum*. For replacement co-culture, *Bl*. *infantis* (or *B. fragilis*) was first grown for 12 hours and then removed by sterile filtration (0.22 µm). The resulting cell-free filtrate (conditioned medium) served as the growth medium for the subsequent inoculation of *Bl*. *longum*. The experimental design is shown in Fig. 7A. In all culturing experiments, each strain was inoculated at an initial OD600 of 0.5 into 400 μL of carbon source-free PYG medium containing 1% 2′-FL as the sole carbon source. Cultures were sampled every 12 hours. At each time point, growth was monitored by measuring the OD600, and an aliquot of the culture was collected and centrifuged for subsequent analysis. All cultivation experiments were performed in triplicate. Bacterial DNA was extracted from the pellet using the DNeasy PowerSoil Pro Kit (Qiagen, Germany) following the manufacturer’s protocol. Quantitative PCR (qPCR) was employed to quantify the relative abundance of each species across all samples. qPCR was performed using a Thermal Cycler Dice Real-Time System (TaKaRa Bio., Japan) with species-specific primer sets, as established in the prior study ^6^. Species-specific quantification was performed using standard curves generated from known concentrations of each species’ genomic DNA.

### Statistical Analysis and Visualization

To investigate which perinatal, lifestyle, and microbiome-related factors associated with the presence or absence of *Bl*. *longum* in Western and Non-western infants, a multivariable logistic regression model was fitted. The logistic regression was performed using the ‘glm’ function in R with a binomial family and logit link. Model estimates, standard errors, Wald statistics, and *p*-values were reported. Datasets with samples taken at less than two months of age were obtained from the COPSAC and Bangladesh cohort containing detailed perinatal, demographic, and metagenomic information of infants at approximately one month of age. The outcome variable was the presence of *Bl*. *longum* in the gut microbiota, represented as a binary variable (0 = absent, 1 = present).

Differences between groups were assessed using non-parametric tests. The Mann-Whitney U test was used for comparisons between two groups, while the Kruskal-Wallis test (a non-parametric alternative to ANOVA) followed by Dunn’s post-hoc test for pairwise comparisons was applied for comparisons among three or more groups. A *p*-value < 0.05 was considered statistically significant.

To compare differences in CAZyme gene composition among different subspecies, we constructed a gene presence/absence matrix for CAZyme genes, calculated the Jaccard distance matrix, and performed non-metric multidimensional scaling (NMDS). Separately, to compare overall genomic gene composition between Western and non-western lifestyle subspecies, we constructed a presence/absence matrix for all genes, calculated the Jaccard distance matrix, and performed principal coordinate analysis (PCoA). To test whether the centroids (i.e., overall composition) differed significantly among the predefined groups (subspecies or lifestyles), we conducted PERMANOVA using the adonis2 function from the vegan package with 999 permutations.

Pearson correlation analysis was conducted for all correlation assessments using the cor.test function in *R* package (v4.5). To compare differences between two independent groups, the Mann–Whitney U test was applied.

Data visualization was carried out using the “boxplot”, “barplot”, “ggplot2”, and “plot” functions in the *R* base package (v4.5).

## RESOURCE AVAILABILITY

### Lead contact

Further information and requests for reagents and resources should be directed to and will be fulfilled by the Lead Contact, Søren J. Sørensen, University of Copenhagen, Urvish Trivedi, University of Copenhagen

## Materials availability

This study did not generate new unique reagents.

## Data and materials availability

One-month metagenomic data from the COPSAC2010 cohort are deposited at the European Nucleotide Archive (ENA) with project accession ID: PRJEB112735. One-year metagenomic sequencing reads have been deposited in the Sequence Read Archive (SRA) under accession number PRJNA715601. Individual-level personally identifiable clinical data from the children participating in the cohort cannot be made publicly available, to protect the privacy of the participants and their families, in accordance with the Danish Data Protection Act and European Regulation 2016/679 of the European Parliament and of the Council (GDPR) that prohibit distribution even in pseudo-anonymized form. However, research collaborations are welcome, and data can be made available under a joint research collaboration by contacting the COPSAC Data Protection Officer (DPO), Ulrik Ralfkiaer, PhD. Requests will be answered within two weeks. Data use is restricted to purposes within childhood health and disease. All non-individual-level data supporting the findings are available from the corresponding author upon reasonable request.

Methodological details regarding all scripts and software packages used in this study are provided in the Methods section.

## Acknowledgements

We gratefully acknowledge the families participating in the COPSAC_2010_ study for their trust and invaluable contributions. We also thank Dr. Olga Sakwinska (Nestlé) for providing metadata from the Bangladesh cohort; Dr. Fredrik Bäckhed (University of Gothenburg) for providing metadata from the Sweden cohort; and Drs. Le Duc Huy Ta and Lee Bee Wah (National University of Singapore) for providing metadata from the Singapore cohort.

This work is supported by the China Scholarship Council, Chengdu Medical College Technology Program (CYYZZ25-15), Independent Research Fund Denmark (grant no. 2032-00103B), Novo Nordisk Foundation [grant no. NNF22OC0073817], and CMC Excellent-talent Program (2024yxGzn05).

COPSAC is funded by private and public research funds all listed at www.copsac.com. The Lundbeck Foundation, The Danish Ministry of Health, Danish Council for Strategic Research, and The Capital Region Research Foundation have provided core support for COPSAC.

## Author contributions

W.G., U.T., and S.J.S. conceived the project. J.S. and J.T. collected the COPSAC2010 samples and associated environmental exposure data. W.G. performed all bioinformatic analyses using partially preprocessed data provided by T.Z. and L.Y. Specifically, T.Z. preprocessed the COPSAC2010 metagenomic data to generate clean reads and genome assemblies. W.Z. and M.D. conducted the *in vitro* culture experiments. X.L. performed DNA extraction from the COPSAC2010 samples. Supervision was provided by U.T. and S.J.S., with key input from T.Z., L.Y., J.S., and J.T. O.S. provided the metadata for the Bangladesh cohort. W.G. drafted the original manuscript and led the writing process, and all authors contributed to manuscript review and editing.

## Reference

1. Laursen, M.F., and Roager, H.M. (2023). Human milk oligosaccharides modify the strength of priority effects in the Bifidobacterium community assembly during infancy. The ISME Journal 17, 2452–2457. 10.1038/s41396-023-01525-7.

2. Olm, M.R., Dahan, D., Carter, M.M., Merrill, B.D., Yu, F.B., Jain, S., Meng, X., Tripathi, S., Wastyk, H., Neff, N., et al. (2022). Robust variation in infant gut microbiome assembly across a spectrum of lifestyles. Science 376, 1220–1223. 10.1126/science.abj2972.

3. Vatanen, T., Ang, Q.Y., Siegwald, L., Sarker, S.A., Le Roy, C.I., Duboux, S., Delannoy-Bruno, O., Ngom-Bru, C., Boulangé, C.L., Stražar, M., et al. (2022). A distinct clade of Bifidobacterium longum in the gut of Bangladeshi children thrives during weaning. Cell 185, 4280–4297.e4212. 10.1016/j.cell.2022.10.011.

4. Barratt, M.J., Nuzhat, S., Ahsan, K., Frese, S.A., Arzamasov, A.A., Sarker, S.A., Islam, M.M., Palit, P., Islam, M.R., Hibberd, M.C., et al. (2022). Bifidobacterium infantis treatment promotes weight gain in Bangladeshi infants with severe acute malnutrition. Sci Transl Med 14, eabk1107. 10.1126/scitranslmed.abk1107.

5. Arzamasov, A.A., Rodionov, D.A., Hibberd, M.C., Guruge, J.L., Kent, J.E., Kazanov, M.D., Leyn, S.A., Elane, M.L., Sejane, K., Furst, A., et al. (2025). Integrative genomic reconstruction reveals heterogeneity in carbohydrate utilization across human gut bifidobacteria. Nat Microbiol. 10.1038/s41564-025-02056-x.

6. Ojima, M.N., Jiang, L., Arzamasov, A.A., Yoshida, K., Odamaki, T., Xiao, J., Nakajima, A., Kitaoka, M., Hirose, J., Urashima, T., et al. (2022). Priority effects shape the structure of infant-type Bifidobacterium communities on human milk oligosaccharides. The ISME Journal 16, 2265–2279. 10.1038/s41396-022-01270-3.

7. O’Callaghan, A., Bottacini, F., O’Connell Motherway, M., and van Sinderen, D. (2015). Pangenome analysis of Bifidobacterium longum and site-directed mutagenesis through by-pass of restriction-modification systems. BMC Genomics 16, 832. 10.1186/s12864-015-1968-4.

8. Shao, Y., Wang, S., Gichuki, B.M., Stares, M.D., Rozday, T.J., Kumar, N., Browne, H.P., Dawson, N.J.R., Njunge, J.M., Tigoi, C., et al. (2026). Genomic atlas of Bifidobacterium infantis and B. longum informs infant probiotic design. Cell 189, 1854–1873.e1817. 10.1016/j.cell.2026.01.007.

9. Modesto, M., Ngom-Bru, C., Scarafile, D., Bruttin, A., Pruvost, S., Sarker, S.A., Ahmed, T., Sakwinska, O., Mattarelli, P., and Duboux, S. (2023). Bifidobacterium longum subsp. iuvenis subsp. nov., a novel subspecies isolated from the faeces of weaning infants. International Journal of Systematic and Evolutionary Microbiology 73. 10.1099/ijsem.0.006013.

10. Vlková, E., Grmanová, M., Killer, J., Mrázek, J., Kopecný, J., Bunesová, V., and Rada, V. (2010). Survival of bifidobacteria administered to calves. Folia Microbiol (Praha) 55, 390–392. 10.1007/s12223-010-0066-x.

11. Sonnenburg, E.D., and Sonnenburg, J.L. (2019). The ancestral and industrialized gut microbiota and implications for human health. Nature Reviews Microbiology 17, 383– 390. 10.1038/s41579-019-0191-8.

12. Dai, D.L.Y., Petersen, C., Hoskinson, C., Del Bel, K.L., Becker, A.B., Moraes, T.J., Mandhane, P.J., Finlay, B.B., Simons, E., Kozyrskyj, A.L., et al. (2023). Breastfeeding enrichment of B. longum subsp. infantis mitigates the effect of antibiotics on the microbiota and childhood asthma risk. Med 4, 92–112.e115. 10.1016/j.medj.2022.12.002.

13. Derrien, M., Mikulic, N., Uyoga, M.A., Chenoll, E., Climent, E., Howard-Varona, A., Nyilima, S., Stoffel, N.U., Karanja, S., Kottler, R., et al. (2023). Gut microbiome function and composition in infants from rural Kenya and association with human milk oligosaccharides. Gut Microbes 15, 2178793. 10.1080/19490976.2023.2178793.

14. Jain, C., Rodriguez-R, L.M., Phillippy, A.M., Konstantinidis, K.T., and Aluru, S. (2018). High throughput ANI analysis of 90K prokaryotic genomes reveals clear species boundaries. Nature Communications 9, 5114. 10.1038/s41467-018-07641-9.

15. Gough, E.K., Edens, T.J., Carr, L., Robertson, R.C., Mutasa, K., Ntozini, R., Chasekwa, B., Geum, H.M., Baharmand, I., Gill, S.K., et al. (2024). Bifidobacterium longum and microbiome maturation modify a nutrient intervention for stunting in Zimbabwean infants. EBioMedicine 108, 105362. 10.1016/j.ebiom.2024.105362.

16. Rasmussen, M., Li, Y., Lindgreen, S., Pedersen, J.S., Albrechtsen, A., Moltke, I., Metspalu, M., Metspalu, E., Kivisild, T., Gupta, R., et al. (2010). Ancient human genome sequence of an extinct Palaeo-Eskimo. Nature 463, 757–762. 10.1038/nature08835.

17. Wibowo, M.C., Yang, Z., Borry, M., Hübner, A., Huang, K.D., Tierney, B.T., Zimmerman, S., Barajas-Olmos, F., Contreras-Cubas, C., García-Ortiz, H., et al. (2021). Reconstruction of ancient microbial genomes from the human gut. Nature 594, 234– 239. 10.1038/s41586-021-03532-0.

18. Willerslev, E., and Meltzer, D.J. (2021). Peopling of the Americas as inferred from ancient genomics. Nature 594, 356–364. 10.1038/s41586-021-03499-y.

19. Moreno-Mayar, J.V., Vinner, L., de Barros Damgaard, P., de la Fuente, C., Chan, J., Spence, J.P., Allentoft, M.E., Vimala, T., Racimo, F., Pinotti, T., et al. (2018). Early human dispersals within the Americas. Science 362. 10.1126/science.aav2621.

20. Potter, B.A., Baichtal, J.F., Beaudoin, A.B., Fehren-Schmitz, L., Haynes, C.V., Holliday, V.T., Holmes, C.E., Ives, J.W., Kelly, R.L., Llamas, B., et al. (2018). Current evidence allows multiple models for the peopling of the Americas. Sci Adv 4, eaat5473. 10.1126/sciadv.aat5473.

21. Ardelean, C.F., Becerra-Valdivia, L., Pedersen, M.W., Schwenninger, J.L., Oviatt, C.G., Macías-Quintero, J.I., Arroyo-Cabrales, J., Sikora, M., Ocampo-Díaz, Y.Z.E., Rubio, C., II, et al. (2020). Evidence of human occupation in Mexico around the Last Glacial Maximum. Nature 584, 87–92. 10.1038/s41586-020-2509-0.

22. Manara, S., Asnicar, F., Beghini, F., Bazzani, D., Cumbo, F., Zolfo, M., Nigro, E., Karcher, N., Manghi, P., Metzger, M.I., et al. (2019). Microbial genomes from non-human primate gut metagenomes expand the primate-associated bacterial tree of life with over 1000 novel species. Genome Biol 20, 299. 10.1186/s13059-019-1923-9.

23. Rühlemann, M.C., Bang, C., Gogarten, J.F., Hermes, B.M., Groussin, M., Waschina, S., Poyet, M., Ulrich, M., Akoua-Koffi, C., Deschner, T., et al. (2024). Functional host- specific adaptation of the intestinal microbiome in hominids. Nature Communications 15, 326. 10.1038/s41467-023-44636-7.

24. Gao, S.-M., Lan, L.-Y., Yang, L., Chen, T., and Fan, P.-F. (2025). Health-associated key gut microbiota drives the variation in community metabolic interactions in non-human primates. Cell Reports 44, 116477. 10.1016/j.celrep.2025.116477.

25. Milani, C., Duranti, S., Napoli, S., Alessandri, G., Mancabelli, L., Anzalone, R., Longhi, G., Viappiani, A., Mangifesta, M., Lugli, G.A., et al. (2019). Colonization of the human gut by bovine bacteria present in Parmesan cheese. Nature Communications 10, 1286. 10.1038/s41467-019-09303-w.

26. Moraïs, S., Winkler, S., Zorea, A., Levin, L., Nagies, F.S.P., Kapust, N., Lamed, E., Artan-Furman, A., Bolam, D.N., Yadav, M.P., et al. (2024). Cryptic diversity of cellulose-degrading gut bacteria in industrialized humans. Science 383, eadj9223. 10.1126/science.adj9223.

27. Kujawska, M., Seki, D., Chalklen, L., Malsom, J., Kiu, R., Goatcher, S., Christoforou, I., Mitra, S., Crouch, L., and Hall, L.J. (2025). Host-specific microbiome and genomic signatures in Bifidobacterium reveal co-evolutionary and functional adaptations across diverse animal hosts. Cell Host Microbe 33, 1502–1517.e1513. 10.1016/j.chom.2025.08.008.

28. Frese, S.A., Benson, A.K., Tannock, G.W., Loach, D.M., Kim, J., Zhang, M., Oh, P.L., Heng, N.C.K., Patil, P.B., Juge, N., et al. (2011). The Evolution of Host Specialization in the Vertebrate Gut Symbiont Lactobacillus reuteri. PLOS Genetics 7, e1001314. 10.1371/journal.pgen.1001314.

29. Trièkoviæ, M., Kieser, S., Zdobnov, E.M., and Trajkovski, M. (2025). Subspecies of the human gut microbiota carry implicit information for in-depth microbiome research. Cell Host & Microbe 33, 1446–1458.e1444. 10.1016/j.chom.2025.07.015.

30. Winglee, K., Howard, A.G., Sha, W., Gharaibeh, R.Z., Liu, J., Jin, D., Fodor, A.A., and Gordon-Larsen, P. (2017). Recent urbanization in China is correlated with a Westernized microbiome encoding increased virulence and antibiotic resistance genes. Microbiome 5, 121 10.1186/s40168-017-0338-7.

31. Odamaki, T., Bottacini, F., Kato, K., Mitsuyama, E., Yoshida, K., Horigome, A., Xiao, J.-z., and van Sinderen, D. (2018). Genomic diversity and distribution of Bifidobacterium longum subsp. longum across the human lifespan. Scientific Reports 8, 85. 10.1038/s41598-017-18391-x.

32. Taft, D.H., Lewis, Z.T., Nguyen, N., Ho, S., Masarweh, C., Dunne-Castagna, V., Tancredi, D.J., Huda, M.N., Stephensen, C.B., Hinde, K., et al. (2022). Bifidobacterium Species Colonization in Infancy: A Global Cross-Sectional Comparison by Population History of Breastfeeding. Nutrients 14. 10.3390/nu14071423.

33. van Best, N., Dominguez-Bello, M.G., Hornef, M.W., Jašarević, E., Korpela, K., and Lawley, T.D. (2022). Should we modulate the neonatal microbiome and what should be the goal? Microbiome 10, 74 10.1186/s40168-022-01281-4.

34. Jarman, J.B., Torres, P.J., Stromberg, S., Sato, H., Stack, C., Ladrillono, A., Pace, S., Jimenez, N.L., Haselbeck, R.J., Insel, R., et al. (2025). Bifidobacterium deficit in United States infants drives prevalent gut dysbiosis. Communications Biology 8, 867. 10.1038/s42003-025-08274-7.

35. Vega-Sagardía, M., Delgado, J., Ruiz-Moyano, S., and Garrido, D. (2023). Proteomic analyses of Bacteroides ovatus and Bifidobacterium longum in xylan bidirectional culture shows sugar cross-feeding interactions. Food Res Int 170, 113025. 10.1016/j.foodres.2023.113025.

36. Sakanaka, M., Gotoh, A., Yoshida, K., Odamaki, T., Koguchi, H., Xiao, J.Z., Kitaoka, M., and Katayama, T. (2019). Varied Pathways of Infant Gut-Associated Bifidobacterium to Assimilate Human Milk Oligosaccharides: Prevalence of the Gene Set and Its Correlation with Bifidobacteria-Rich Microbiota Formation. Nutrients 12. 10.3390/nu12010071.

37. Marcobal, A., Barboza, M., Sonnenburg, Erica D., Pudlo, N., Martens, Eric C., Desai, P., Lebrilla, Carlito B., Weimer, Bart C., Mills, David A., German, J.B., and Sonnenburg, Justin L. (2011). Bacteroides in the Infant Gut Consume Milk Oligosaccharides via Mucus-Utilization Pathways. Cell Host & Microbe 10, 507–514. 10.1016/j.chom.2011.10.007.

38. Bäckhed, F., Roswall, J., Peng, Y., Feng, Q., Jia, H., Kovatcheva-Datchary, P., Li, Y., Xia, Y., Xie, H., Zhong, H., et al. (2015). Dynamics and Stabilization of the Human Gut Microbiome during the First Year of Life. Cell Host Microbe 17, 690–703. 10.1016/j.chom.2015.04.004.

39. Li, X., Stokholm, J., Brejnrod, A., Vestergaard, G.A., Russel, J., Trivedi, U., Thorsen, J., Gupta, S., Hjelmsø, M.H., Shah, S.A., et al. (2021). The infant gut resistome associates with E. coli, environmental exposures, gut microbiome maturity, and asthma-associated bacterial composition. Cell Host Microbe 29, 975–987.e974. 10.1016/j.chom.2021.03.017.

40. Robertson, R.C., Edens, T.J., Carr, L., Mutasa, K., Gough, E.K., Evans, C., Geum, H.M., Baharmand, I., Gill, S.K., Ntozini, R., et al. (2023). The gut microbiome and early-life growth in a population with high prevalence of stunting. Nature Communications 14, 654. 10.1038/s41467-023-36135-6.

41. D’Souza, A.W., Moodley-Govender, E., Berla, B., Kelkar, T., Wang, B., Sun, X., Daniels, B., Coutsoudis, A., Trehan, I., and Dantas, G. (2020). Cotrimoxazole Prophylaxis Increases Resistance Gene Prevalence and á-Diversity but Decreases β-Diversity in the Gut Microbiome of Human Immunodeficiency Virus-Exposed, Uninfected Infants. Clin Infect Dis 71, 2858–2868. 10.1093/cid/ciz1186.

42. Kim, M., Rodriguez-R, L.M., Hatt, J.K., Kayali, O., Nalá, R., Dunlop, A.L., Brennan, P.A., Corwin, E., Smith, A.K., Brown, J., and Konstantinidis, K.T. (2022). Higher pathogen load in children from Mozambique vs. USA revealed by comparative fecal microbiome profiling. ISME Communications 2, 74. 10.1038/s43705-022-00154-z.

43. Tett, A., Huang, K.D., Asnicar, F., Fehlner-Peach, H., Pasolli, E., Karcher, N., Armanini, F., Manghi, P., Bonham, K., Zolfo, M., et al. (2019). The Prevotella copri Complex Comprises Four Distinct Clades Underrepresented in Westernized Populations. Cell Host Microbe 26, 666–679.e667. 10.1016/j.chom.2019.08.018.

44. Vatanen, T., Jabbar, K.S., Ruohtula, T., Honkanen, J., Avila-Pacheco, J., Siljander, H., Stražar, M., Oikarinen, S., Hyöty, H., Ilonen, J., et al. (2022). Mobile genetic elements from the maternal microbiome shape infant gut microbial assembly and metabolism. Cell 185, 4921–4936.e4915. 10.1016/j.cell.2022.11.023.

45. Shao, Y., Forster, S.C., Tsaliki, E., Vervier, K., Strang, A., Simpson, N., Kumar, N., Stares, M.D., Rodger, A., Brocklehurst, P., et al. (2019). Stunted microbiota and opportunistic pathogen colonization in caesarean-section birth. Nature 574, 117– 121. 10.1038/s41586-019-1560-1.

46. Murphy, R., Morgan, X.C., Wang, X.Y., Wickens, K., Purdie, G., Fitzharris, P., Otal, A., Lawley, B., Stanley, T., Barthow, C., et al. (2019). Eczema-protective probiotic alters infant gut microbiome functional capacity but not composition: sub-sample analysis from a RCT. Benef Microbes 10, 5–17. 10.3920/bm2017.0191.

47. Ta, L.D.H., Chan, J.C.Y., Yap, G.C., Purbojati, R.W., Drautz-Moses, D.I., Koh, Y.M., Tay, C.J.X., Huang, C.H., Kioh, D.Y.Q., Woon, J.Y., et al. (2020). A compromised developmental trajectory of the infant gut microbiome and metabolome in atopic eczema. Gut Microbes 12, 1–22. 10.1080/19490976.2020.1801964.

48. Ferretti, P., Pasolli, E., Tett, A., Asnicar, F., Gorfer, V., Fedi, S., Armanini, F., Truong, D.T., Manara, S., Zolfo, M., et al. (2018). Mother-to-Infant Microbial Transmission from Different Body Sites Shapes the Developing Infant Gut Microbiome. Cell Host Microbe 24, 133–145.e135. 10.1016/j.chom.2018.06.005.

49. Manara, S., Selma-Royo, M., Huang, K.D., Asnicar, F., Armanini, F., Blanco-Miguez, A., Cumbo, F., Golzato, D., Manghi, P., Pinto, F., et al. (2023). Maternal and food microbial sources shape the infant microbiome of a rural Ethiopian population. Curr Biol 33, 1939–1950.e1934. 10.1016/j.cub.2023.04.011.

50. Cait, A., Cardenas, E., Dimitriu, P.A., Amenyogbe, N., Dai, D., Cait, J., Sbihi, H., Stiemsma, L., Subbarao, P., Mandhane, P.J., et al. (2019). Reduced genetic potential for butyrate fermentation in the gut microbiome of infants who develop allergic sensitization. J Allergy Clin Immunol 144, 1638–1647.e1633. 10.1016/j.jaci.2019.06.029.

51. Baumann-Dudenhoeffer, A.M., D’Souza, A.W., Tarr, P.I., Warner, B.B., and Dantas, G. (2018). Infant diet and maternal gestational weight gain predict early metabolic maturation of gut microbiomes. Nat Med 24, 1822–1829. 10.1038/s41591-018-0216-2.

52. Casaburi, G., Duar, R.M., Brown, H., Mitchell, R.D., Kazi, S., Chew, S., Cagney, O., Flannery, R.L., Sylvester, K.G., Frese, S.A., et al. (2021). Metagenomic insights of the infant microbiome community structure and function across multiple sites in the United States. Scientific Reports 11, 1472. 10.1038/s41598-020-80583-9.

53. Xiao, L., Wang, J., Zheng, J., Li, X., and Zhao, F. (2021). Deterministic transition of enterotypes shapes the infant gut microbiome at an early age. Genome Biol 22, 243. 10.1186/s13059-021-02463-3.

54. Joshi NA, F.J. (2011). Sickle: A sliding-window, adaptive,quality-based trimming tool for FastQ files.

55. Bushnell, B. (2015). BBMap short-read aligner, and other bioinformatics tools.

56. Nurk, S., Meleshko, D., Korobeynikov, A., and Pevzner, P.A. (2017). metaSPAdes: a new versatile metagenomic assembler. Genome Res 27, 824–834. 10.1101/gr.213959.116.

57. Uritskiy, G.V., DiRuggiero, J., and Taylor, J. (2018). MetaWRAP-a flexible pipeline for genome-resolved metagenomic data analysis. Microbiome 6, 158. 10.1186/s40168-018-0541-1.

58. Nissen, J.N., Johansen, J., Allesøe, R.L., Sønderby, C.K., Armenteros, J.J.A., Grønbech, C.H., Jensen, L.J., Nielsen, H.B., Petersen, T.N., Winther, O., and Rasmussen, S. (2021). Improved metagenome binning and assembly using deep variational autoencoders. Nature Biotechnology 39, 555–560. 10.1038/s41587-020-00777-4.

59. Parks, D.H., Imelfort, M., Skennerton, C.T., Hugenholtz, P., and Tyson, G.W. (2015). CheckM: assessing the quality of microbial genomes recovered from isolates, single cells, and metagenomes. Genome Res 25, 1043–1055. 10.1101/gr.186072.114.

60. Chaumeil, P.A., Mussig, A.J., Hugenholtz, P., and Parks, D.H. (2019). GTDB-Tk: a toolkit to classify genomes with the Genome Taxonomy Database. Bioinformatics 36, 1925–1927. 10.1093/bioinformatics/btz848.

61. Olm, M.R., Brown, C.T., Brooks, B., and Banfield, J.F. (2017). dRep: a tool for fast and accurate genomic comparisons that enables improved genome recovery from metagenomes through de-replication. The ISME Journal 11, 2864–2868. 10.1038/ismej.2017.126.

62. Segata, N., Börnigen, D., Morgan, X.C., and Huttenhower, C. (2013). PhyloPhlAn is a new method for improved phylogenetic and taxonomic placement of microbes. Nature Communications 4, 2304. 10.1038/ncomms3304.

63. Buchfink, B., Xie, C., and Huson, D.H. (2015). Fast and sensitive protein alignment using DIAMOND. Nature Methods 12, 59–60. 10.1038/nmeth.3176.

64. Katoh, K., and Standley, D.M. (2013). MAFFT multiple sequence alignment software version 7: improvements in performance and usability. Mol Biol Evol 30, 772–780. 10.1093/molbev/mst010.

65. Capella-Gutiérrez, S., Silla-Martínez, J.M., and Gabaldón, T. (2009). trimAl: a tool for automated alignment trimming in large-scale phylogenetic analyses. Bioinformatics 25, 1972–1973. 10.1093/bioinformatics/btp348.

66. Price, M.N., Dehal, P.S., and Arkin, A.P. (2010). FastTree 2--approximately maximum-likelihood trees for large alignments. PLoS One 5, e9490. 10.1371/journal.pone.0009490.

67. Stamatakis, A. (2014). RAxML version 8: a tool for phylogenetic analysis and post-analysis of large phylogenies. Bioinformatics 30, 1312–1313. 10.1093/bioinformatics/btu033.

68. Page, A.J., Cummins, C.A., Hunt, M., Wong, V.K., Reuter, S., Holden, M.T., Fookes, M., Falush, D., Keane, J.A., and Parkhill, J. (2015). Roary: rapid large-scale prokaryote pan genome analysis. Bioinformatics 31, 3691–3693. 10.1093/bioinformatics/btv421.

69. Yu, G., Smith, D.K., Zhu, H., Guan, Y., and Lam, T.T.-Y. (2017). ggtree: an r package for visualization and annotation of phylogenetic trees with their covariates and other associated data. Methods in Ecology and Evolution 8, 28–36. 10.1111/2041-210X.12628.

70. Asnicar, F., Weingart, G., Tickle, T.L., Huttenhower, C., and Segata, N. (2015). Compact graphical representation of phylogenetic data and metadata with GraPhlAn. PeerJ 3, e1029. 10.7717/peerj.1029.

71. Bouckaert, R., Vaughan, T.G., Barido-Sottani, J., Duchêne, S., Fourment, M., Gavryushkina, A., Heled, J., Jones, G., Kühnert, D., De Maio, N., et al. (2019). BEAST 2.5: An advanced software platform for Bayesian evolutionary analysis. PLoS Comput Biol 15, e1006650. 10.1371/journal.pcbi.1006650.

72. Croucher, N.J., Page, A.J., Connor, T.R., Delaney, A.J., Keane, J.A., Bentley, S.D., Parkhill, J., and Harris, S.R. (2015). Rapid phylogenetic analysis of large samples of recombinant bacterial whole genome sequences using Gubbins. Nucleic Acids Res 43, e15. 10.1093/nar/gku1196.

73. Drummond, A.J., Ho, S.Y., Phillips, M.J., and Rambaut, A. (2006). Relaxed phylogenetics and dating with confidence. PLoS Biol 4, e88. 10.1371/journal.pbio.0040088.

74. Rambaut, A., Drummond, A.J., Xie, D., Baele, G., and Suchard, M.A. (2018). Posterior Summarization in Bayesian Phylogenetics Using Tracer 1.7. Syst Biol 67, 901–904. 10.1093/sysbio/syy032.

75. Pritchard, L., Glover, R.H., Humphris, S., Elphinstone, J.G., and Toth, I.K.J.A.m. (2016). Genomics and taxonomy in diagnostics for food security: soft-rotting enterobacterial plant pathogens. 8, 12–24. 10.1039/c5ay02550h.

76. Löytynoja, A., Vilella, A.J., and Goldman, N.J.B. (2012). Accurate extension of multiple sequence alignments using a phylogeny-aware graph algorithm. 28, 1684–1691.

77. Hyatt, D., Chen, G.-L., LoCascio, P.F., Land, M.L., Larimer, F.W., and Hauser, L.J. (2010). Prodigal: prokaryotic gene recognition and translation initiation site identification. BMC Bioinformatics 11, 119. 10.1186/1471-2105-11-119.

78. Langmead, B., and Salzberg, S.L. (2012). Fast gapped-read alignment with Bowtie 2. Nature Methods 9, 357–359. 10.1038/nmeth.1923.

79. Olm, M.R., Crits-Christoph, A., Bouma-Gregson, K., Firek, B.A., Morowitz, M.J., and Banfield, J.F. (2021). inStrain profiles population microdiversity from metagenomic data and sensitively detects shared microbial strains. Nature Biotechnology 39, 727–736. 10.1038/s41587-020-00797-0.

80. Fu, L., Niu, B., Zhu, Z., Wu, S., and Li, W. (2012). CD-HIT: accelerated for clustering the next-generation sequencing data. Bioinformatics 28, 3150–3152. 10.1093/bioinformatics/bts565.

81. Li, H., and Durbin, R. (2009). Fast and accurate short read alignment with Burrows-Wheeler transform. Bioinformatics 25, 1754–1760. 10.1093/bioinformatics/btp324.

82. Kang, D.D., Li, F., Kirton, E., Thomas, A., Egan, R., An, H., and Wang, Z. (2019). MetaBAT 2: an adaptive binning algorithm for robust and efficient genome reconstruction from metagenome assemblies. PeerJ 7, e7359. 10.7717/peerj.7359.

83. Zheng, J., Ge, Q., Yan, Y., Zhang, X., Huang, L., and Yin, Y. (2023). dbCAN3: automated carbohydrate-active enzyme and substrate annotation. Nucleic Acids Research 51, W115–W121. 10.1093/nar/gkad328.

84. Lombard, V., Golaconda Ramulu, H., Drula, E., Coutinho, P.M., and Henrissat, B. (2014). The carbohydrate-active enzymes database (CAZy) in 2013. Nucleic Acids Res 42, D490–495. 10.1093/nar/gkt1178.

85. Gilchrist, C.L.M., and Chooi, Y.-H. (2021). clinker &amp; clustermap.js: automatic generation of gene cluster comparison figures. Bioinformatics 37, 2473–2475. 10.1093/bioinformatics/btab007.

86. Aziz, R.K., Bartels, D., Best, A.A., DeJongh, M., Disz, T., Edwards, R.A., Formsma, K., Gerdes, S., Glass, E.M., Kubal, M., et al. (2008). The RAST Server: rapid annotations using subsystems technology. BMC Genomics 9, 75. 10.1186/1471-2164-9-75.

87. Teufel, F., Almagro Armenteros, J.J., Johansen, A.R., Gíslason, M.H., Pihl, S.I., Tsirigos, K.D., Winther, O., Brunak, S., von Heijne, G., and Nielsen, H. (2022). SignalP 6.0 predicts all five types of signal peptides using protein language models. Nat Biotechnol 40, 1023–1025. 10.1038/s41587-021-01156-3.

