## Supplementary Figures for "Evolutionary history and microbial cross-feeding shape lifestyle-stratified dominance of *Bifidobacterium longum* subspecies in infants"

KEY RESOURCES TABLE

| REAGENT or RESOURCE | SOURCE | IDENTIFIER |
|---|---|---|
| Biological samples | | |
| Infant feces samples | This study | COPSAC2010 cohort |
| Deposited infant metagenome raw data | | |
| Bangladesh | Vatanen et al. | PRJNA806984 |
| Zimbabwe | Robertson et al. | PRJEB51728 |
| Kenya | Derrien et al. | PRJEB52748 |
| South Africa | D'Souza et al. | PRJNA549787 |
| Mozambique/USA | Kim et al. | PRJNA747761 |
| Ethiopia | Tett et al. | PRJNA504891 |
| Tanzania (Hadza) | Olm et al. | PRJEB49206 |
| Sweden | Backhed et al. | PRJEB6456 |
| Finland | Vatanen et al. | PRJNA821542 |
| UK | Shao et al. | PRJEB32631 |
| New Zealand | Murphy et al. | PRJNA345144 |
| Singapore | Ta et al. | PRJNA642723 |
| Italy | Ferretti et al. | PRJNA352475 |
| Italy | Manara et al. | PRJNA716780 |
| Canada | Cait et al. | PRJEB32135 |
| USA | Casaburi et al | PRJNA633576 |
| USA | Aimee et al. | PRJNA473126 |
| China | Xiao et al. | PRJNA695070 |
| Deposited cattle metagenome raw data | | |
| Cattle (China) | Zhuang et al. | PRJNA1052964 |
| Cattle (China) | Gao et al. | PRJNA1042798 |
| Cattle (USA) | Salaheen et al. | PRJNA645414 |
| Cattle (USA) | Drake et al. | PRJNA1088536 |
| Deposited NHP metagenome raw data | | |

| | | |
|---|---|---|
| Gorilla gorilla | Amato et al.; Sharma et al; Campbell et al | ERP104379; PRJNA635116; PRJNA539933 |
| Pan troglodytes | Amato et al.; Campbell et al.; Sanders et al. | ERP104379; PRJNA539933; PRJNA842587 |
| Gorilla gorilla gorilla | Hicks et al. | PRJNA382701 |
| Papio cynocephalus | Tung et al. | PRJNA271618 |
| Macaca fascicularis | Li et al.; Duanet al. | PRJEB22765; SRP218382 |
| Cebus capucinus imitator | Orkin et al. | PRJNA485217 |
| Trachypithecus leucocephalus | Que et al. | CRA002281 |
| Macaca fuscata | Hanya et al. | DRA009275 |
| Rhinopithecus bieti | Xia et al. | PRJNA787384 |
| Rhinopithecus roxellana | Li et al.; Li et al.; Zhang et al. | PRJNA799478 |
| Macaca mulatta | Li et al.; Rhoades et al.; Nicholas et al. | PRJNA799478; PRJNA862694; PRJNA954012; PRJNA896946; PRJNA1017507; PRJNA546004 |
| Propithecus coquereli | Greene et al. | PRJNA495032; PRJNA684050 |
| Hoolock tianxing | Li et al. | PRJCA012504 |
| Nomascus hainanus | Huang et al. | CRA019990 |
| Nomascus concolor | Gao et al. | PRJNA1158499 |
| Deposited Ancient human metagenome raw data | | |
| Palaeofaeces (Mexico) | Hagan et al. | PRJEB33577 |
| Digestive tract contents (Italy) | Maixner et al. | PRJEB11511 |
| Palaeofaeces (Austria) | Maixner et al. | PRJEB44507 |
| Palaeofaeces (Mexico) | Tett et al. | PRJEB31971 |
| Palaeofaeces (Mexico; USA) | Wibowo et al. | PRJNA561510 |
| Software and algorithms | | |
| BBDuk v39.01 | https://sourceforge.net/projects/bbmap/ | https://jgi.doe.gov/data-and-tools/bbtools |
| Sickle v1.33 | Joshi and Fass, 2011 | https://github.com/najoshi/sickle/releases |
| BBMap v38.9 | Bushnell et al. 2015 | https://github.com/bbushnell/BBTools |
| SPAdes v3.15.5 | Nurk et al., 2017 | https://cab.spbu.ru/files/release3.12.0/manual.html |
| MetaWRAP v1.3.2 | Uritskiy et al., 2018 | https://github.com/bxlab/metaWRAP |
| VAMB v3.0.8 | Nissen et al., 2021 | https://github.com/RasmussenLab/vamb |
| MaxBin2 v2.2.7 | Wu et al., 2016 | https://sourceforge.net/projects/maxbin2/files |
| MetaBAT2 v2.13 | Kang et al., 2019 | https://bitbucket.org/berkeleylab/metabat |
| CONCOCT v0.4 | Alneberg et al., 2014 | https://github.com/BinPro/CONCOCT |
| CheckM v1.2.2 | Parks et al., 2015 | https://github.com/Ecogenomics/CheckM |

| | | |
|---|---|---|
| GTDB-tk v2.1 | Parks et al., 2018 | https://github.com/Ecogenomics/GTDBTk |
| dRep v3.4.3 | Olm et al., 2017 | https://github.com/MrOlm/drep |
| PhyloPhlAn v3.0.68 | Segata et al., 2013 | https://github.com/biobakery/phylophlan |
| DIAMOND v2.0.13 | Buchfink et al., 2015 | https://github.com/bbuchfink/diamond/releases |
| MAFFT v7.490 | Katoh et al., 2013 | https://mafft.cbrc.jp/alignment/software/source.html |
| trimAl v1.4.1 | Capella-Gutiérrez et al., 2009 | https://trimal.readthedocs.io/en/latest |
| FastTree v2.1.11 | Price et la., 2010 | http://bioconda.github.io/recipes/fasttree |
| RaxML v8.2.12 | Stamatakis et al., 2014 | https://evomics.org/learning/phylogenetics/raxml |
| GraPhlAn v1.13 | Asnicar et al., 2015 | https://huttenhower.sph.harvard.edu/graphlan |
| R v4.3.3 | core Team, 2018 | https://www.r-project.org |
| pyani v0.2.11 | Gower et al., 2020 | https://github.com/widdowquinn/pyani |
| Roary v3.13.0 | Page et al., 2015 | https://github.com/sanger-pathogens/Roary |
| Bowtie2 v2.3.5 | Langmead et la., 2012 | https://bowtie-bio.sourceforge.net/bowtie2 |
| inStrain vv1.3.4 | Olm et al., 2021 | https://github.com/MrOlm/inStrain |
| CD-HIT v4.8.1 | Fu et al., 2012 | https://www.bioinformatics.org/cd-hit |
| BWA v0.1.79 | Li et al., 2009 | https://github.com/lh3/bwa |
| SignalP v6.0 | Teufel et al., 2020 | http://www.cbs.dtu.dk/services/SignalP |
| BEAST v2.7.7 | Bouckaert *et al.*, 2019 | https://beast.community/ |
| Gubbins v3.4.3 | Croucher *et al.*, 2015 | https://github.com/nickjcroucher/gubbins |
| dbCAN v3 | Zheng et al., 2023 | https://pro.unl.edu/dbCAN2 |

### Supplementary Figures and Tables

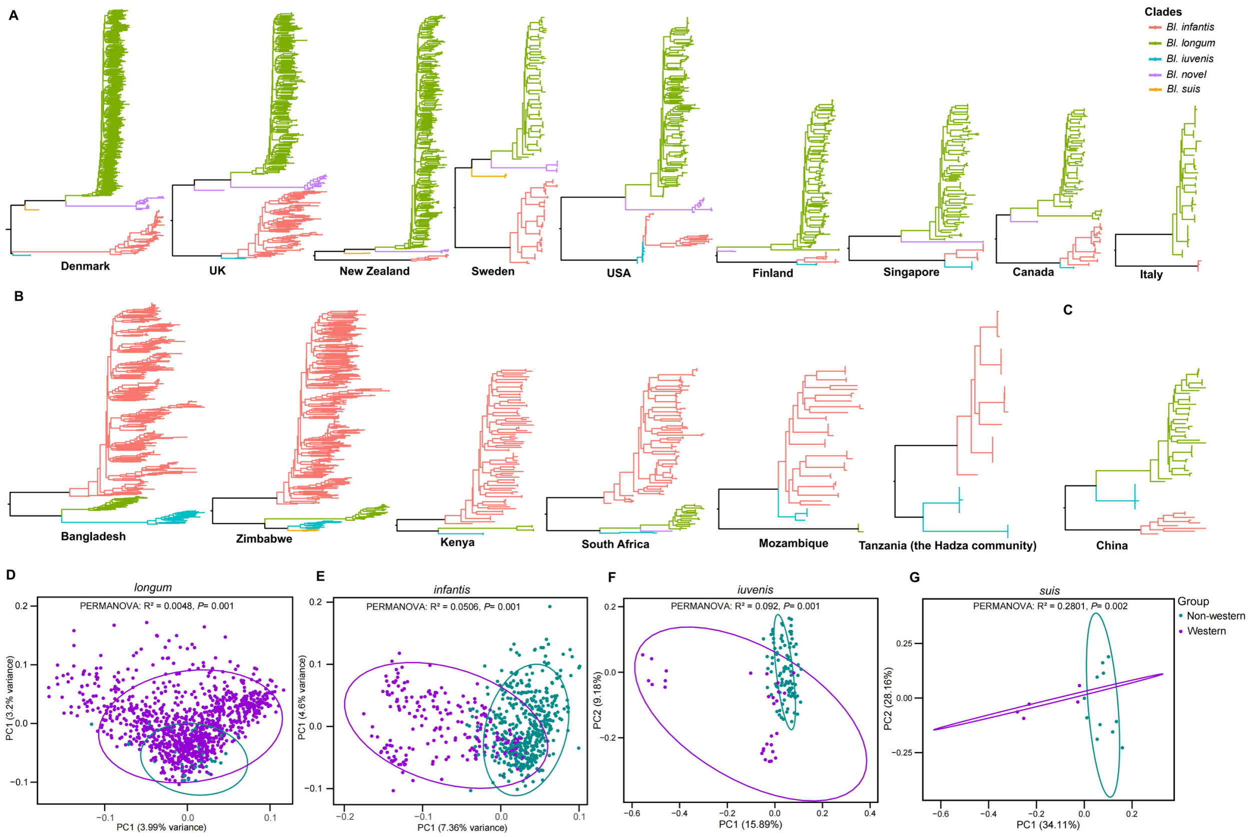


**Fig. S1. Phylogenetic tree and PCoA (based on gene presence–absence) of infant gut *B. longum* subspecies from Western and non-Western lifestyles.** Phylogenetic tree of infant gut-derived *B. longum* from **(A)** Western and **(B)** Non-Western lifestyles; **(C)** Phylogenetic tree of infant gut-derived *B. longum* from China; PCoA of gene presence/absence using Jaccard distances for **(D)** *Bl. longum*, **(E)** *Bl. infantis*, **(F)** *Bl. iuvenis*, and **(G)** *Bl. suis* Variance explained by lifestyle was calculated by PERMANOVA calculated with 999 permutations.

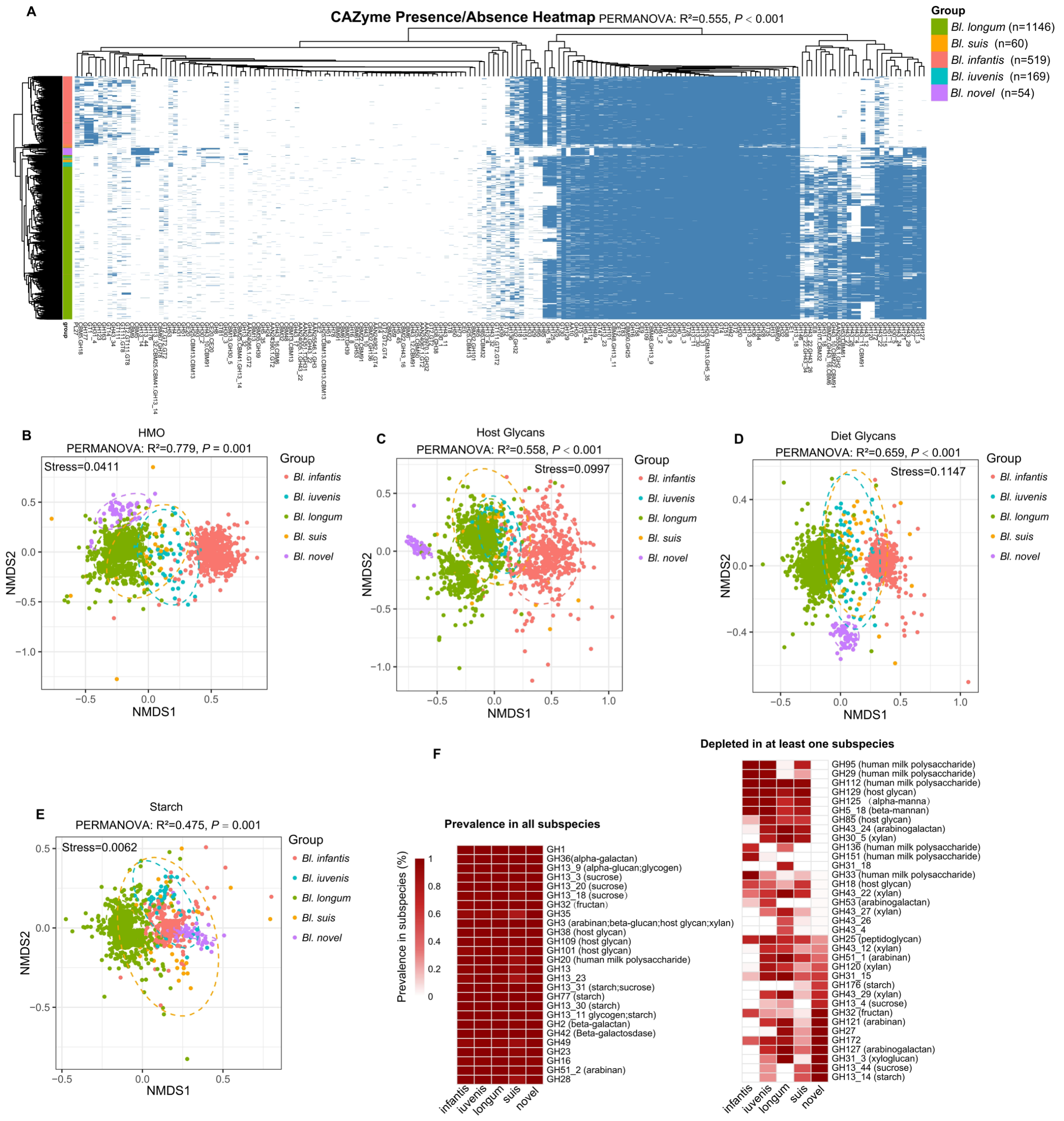


**Fig. S2. Functional Diversity of the *B. longum* subspecies. (A)** Heatmap showing the presence (blue) or absence (white) of genes encoding CAZy functions across the five *B. longum* subspecies; **(B)** NMDS ordination using Bray-Curtis distances of CAZy families involved in HMO, **(C)** host glycans, **(D)** dietary glycans, and **(E)** starch utilization across different *B. longum* subspecies; **(F)** CAZy families significantly enriched (left; prevalence > 80% in all subspecies) or depleted (right; in at least one subspecies compared to the other subspecies).

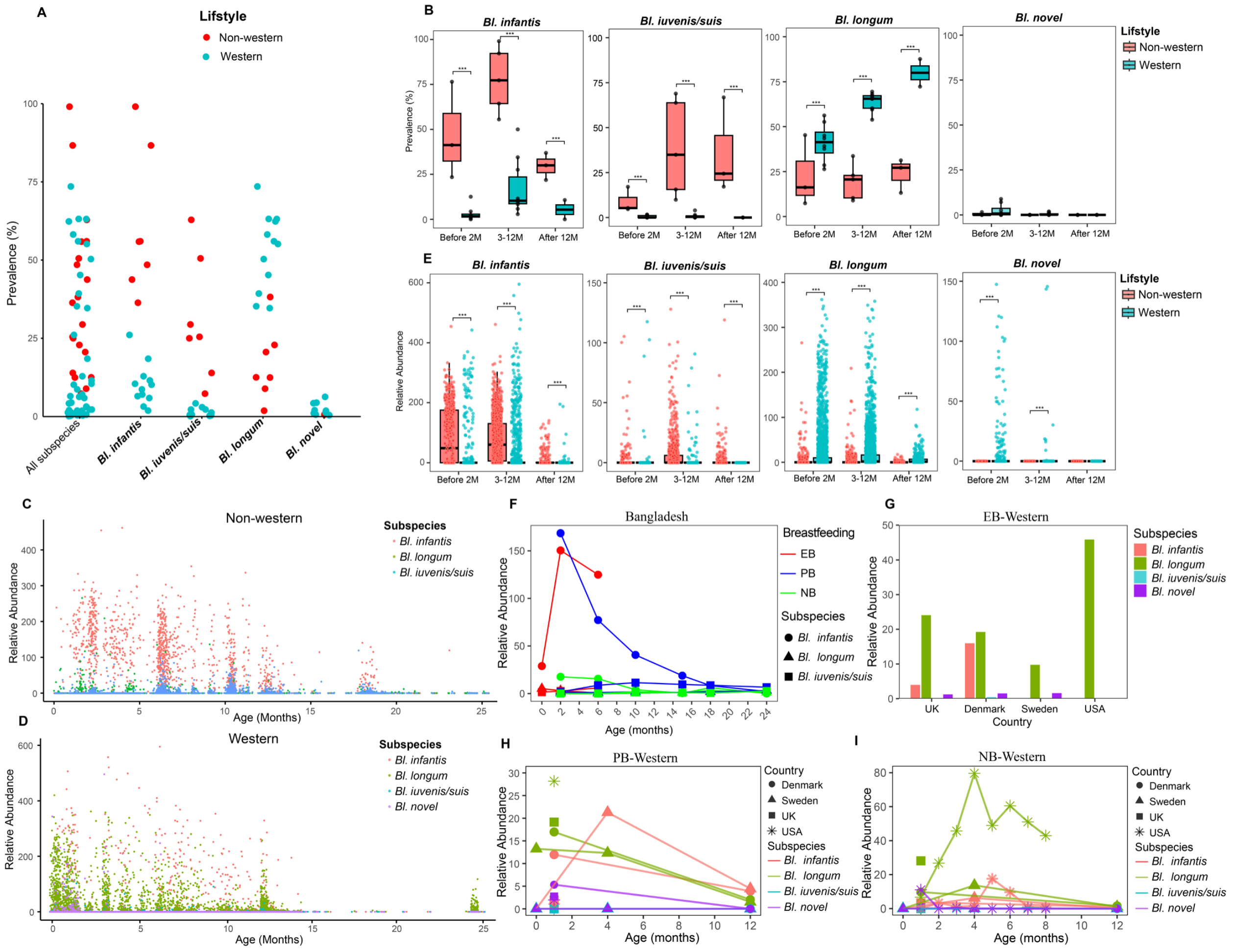


**Fig. S3. Prevalence and relative abundance of *B. longum* subspecies in infants from non-Westernized and Westernized populations. (A)** *B. longum* prevalence in non-Westernized and Westernized datasets. Each dot is the average prevalence from a dataset. "All subspecies" refers to the prevalence of any of the subspecies being present; **(B)** Box plots showing the prevalence and relative abundance **(E)** of *Bl. infantis*, *Bl. iuvenis/suis*, *Bl. longum* (BT), and *Bl. novel* across different age groups (Before 2 months, 3-12 months, and after 12 months) in non-Westernized and Westernized infants; *B. longum* subspecies relative abundance across infant ages in non-western **(C)** and western **(D)** populations; **(F)** Abundance dynamics of *B. longum* subspecies in Bangladesh infants by breastfeeding pattern; Abundance dynamics of *B. longum* subspecies in western lifestyle infants with exclusive breastfeeding **(G)**, partial breastfeeding **(H)** and no breast feeding **(I).**

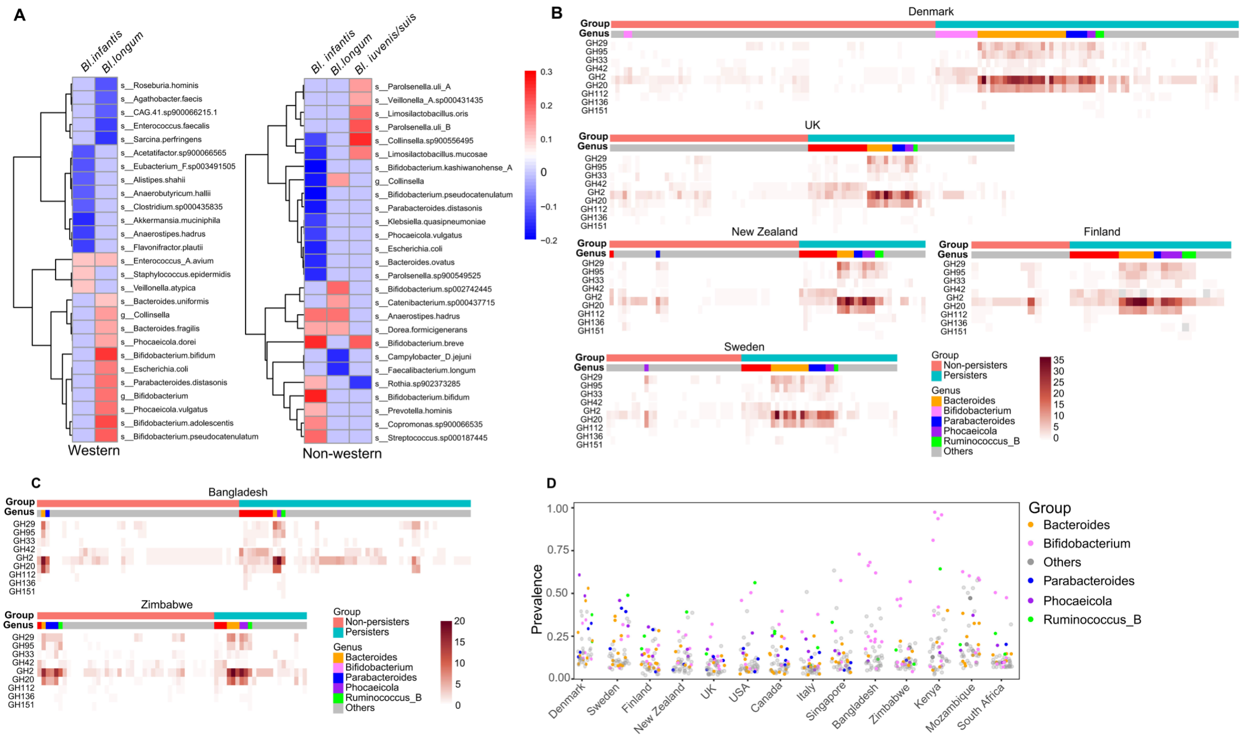

**Fig. S4. Characteristics of persistent gut strains from Western and non-Western infants. (A)** Heatmap of correlations between persistence of *B. longum* toother strains; Heatmaps of CAZy genes related to human milk oligosaccharide (HMO) degradation in persistent versus non-persistent strains fromWestern **(B)** and non-Western **(C)** infants; (**D)** Top 50 most prevalent bacterial strains harboring GH29, GH95, and GH33 glycoside hydrolase genes across geographic cohorts.

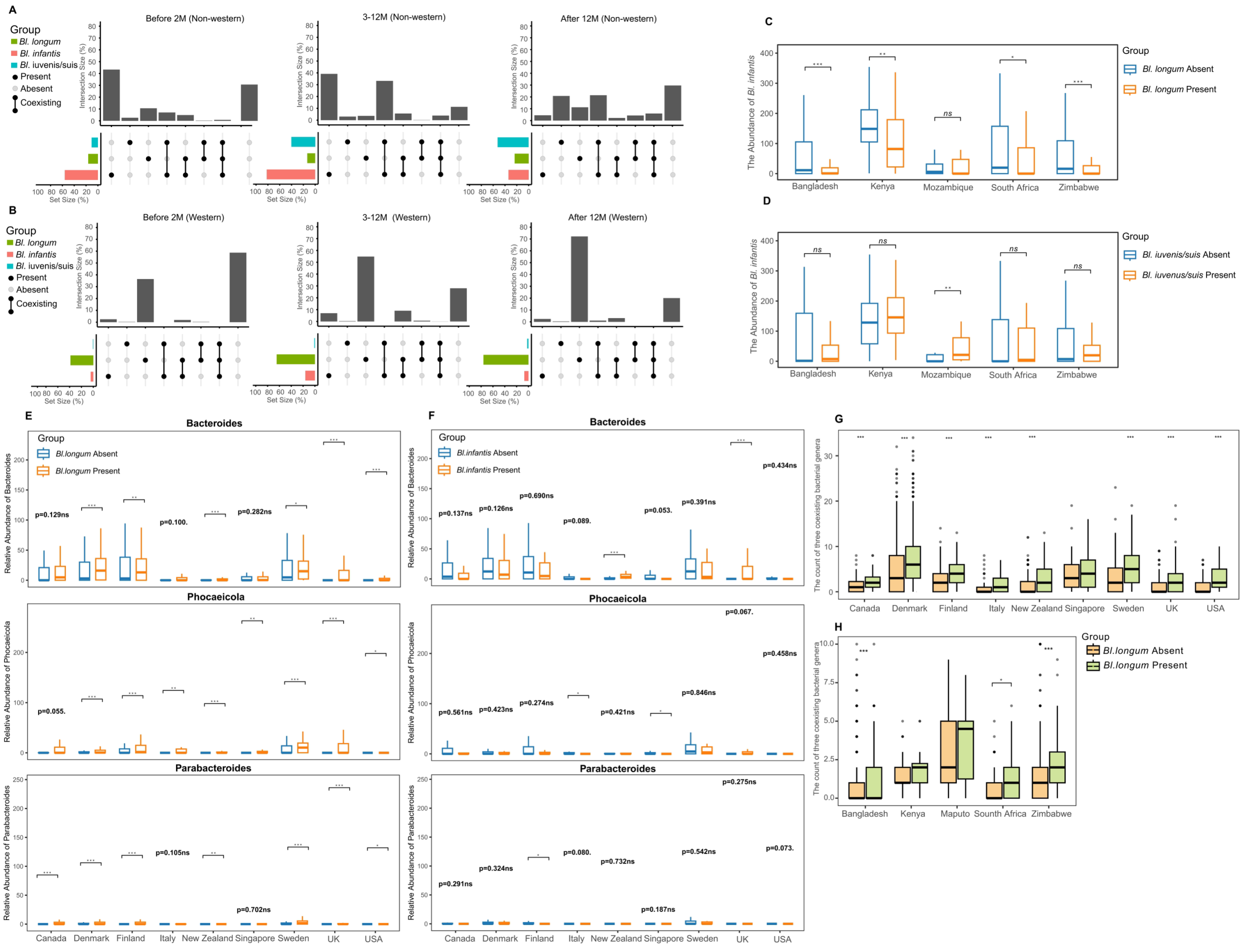


**Fig. S5. Subspecies prevalence, interspecific co-occurrence, and relative abundance/species count of *Bacteroides*, *Phocaeicola*, and *Parabacteroides* in infants with *B. longum* subsp. *longum* or *infantis* colonization.** UPset plots display the prevalence and co-occurrence patterns of *B. longum* subspecies in **(A)** non-Westernized and **(B)** Westernized infants across the same age intervals; Assessing the relative abundance of *Bl. infantis* in relation to *Bl. longum* presence vs. *Bl. longum* absence **(C)**, and *Bl. iuvenis/suis* presence vs. *Bl. iuvenis/suis* absence **(D)** in the gut of infants with western lifestyle; Assessing the relative abundance of *Bacteroides*, *Phocaeicola*, and *Parabacteroides* in relation to *Bl. longum* presence vs. *Bl. longum* absence **(E)**, and *Bl. infantis* presence vs. *Bl. infantis* absence **(F)** in the gut of infants western lifestyle; Comparison of the number of species within the genera Bacteroides, Phocaeicola, and Parabacteroides between infants harboring and those lacking *Bl. longum* in countries with **(G)** Western and **(H)** non-Western lifestyle. * $p < 0.05$; ** $p < 0.01$; *** $p < 0.001$.

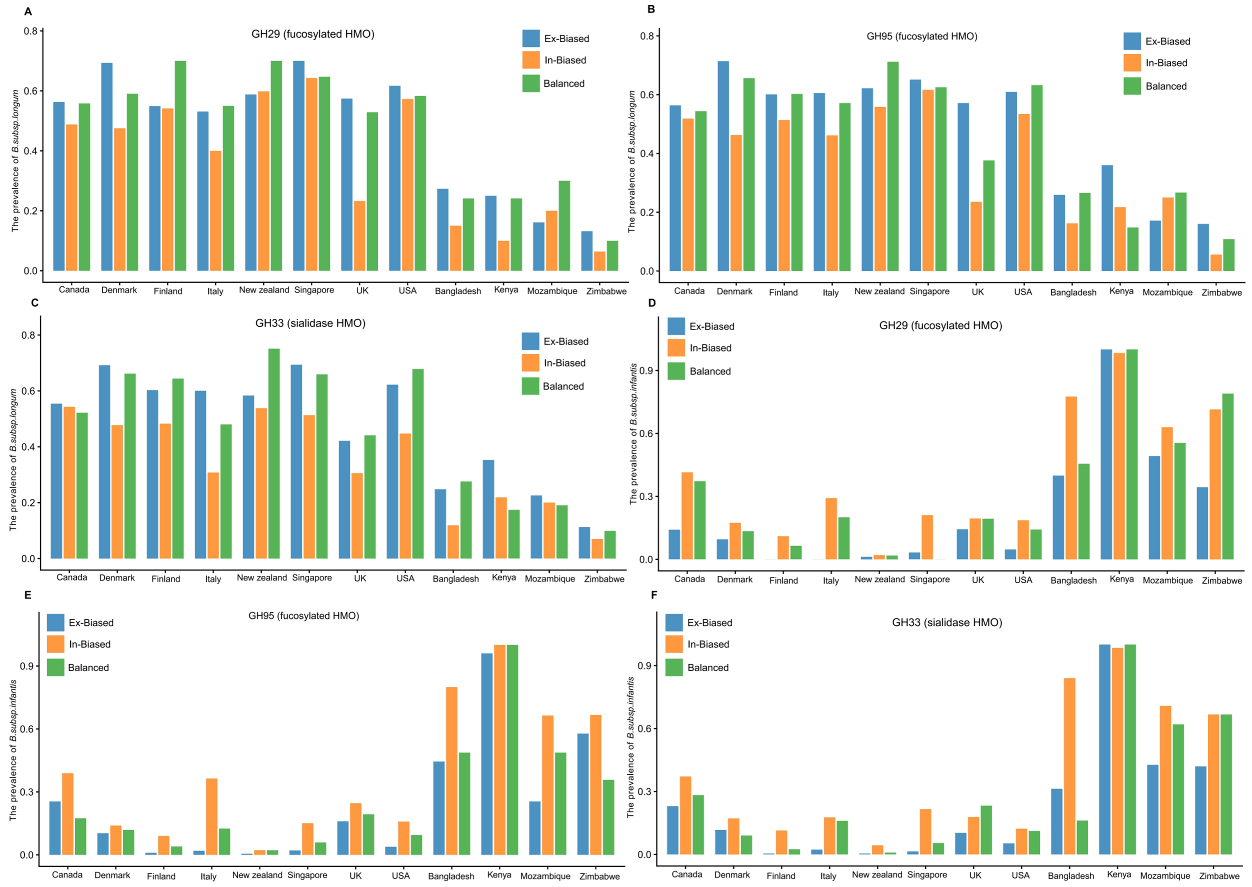


**Fig S6. Prevalence of *B. longum* subsp. *longum* and *B. longum* subsp. *infantis*, Stratified by Gene Group (Ex-dominant, In-dominant, Intermediate). (A–C)** Detection rates of *B. longum* subsp. *longum*, stratified by gene (GH29, GH95, GH33) and subsequently categorized into Ex-dominant, In-dominant, or Intermediate groups based on the extracellular vs. intracellular gene abundance ratio. **(D–F)** Detection rates of *B. longum* subsp. *infantis*, stratified by gene (GH29, GH95, GH33) and subsequently categorized into Ex-dominant, In-dominant, or Intermediate groups based on the extracellular vs. intracellular gene abundance ratio. Ex-dominant: $\log_2$ (Extracellular abundance/Intracellular abundance) $\geq 1$; In-dominant: $\log_2$ (Extracellular abundance/Intracellular abundance) $\leq -1$; Intermediate: $-1 < \log_2$ (Extracellular abundance / Intracellular abundance) $< 1$.

**Table S1. Details of assembled high-quality MAGs in this study.**

**Table S2. Summary of reconstructed *B. longum* bins and subspecies counts by country.**

**Table S3. List of MAGs and genomes employed in the construction of the phylogenetic tree of *Bl. longum* or *Bl. iuvenis/suis*.**

**Table S4. Prevalence of HMO Utilization Genes Across *B. longum* Subspecies.**

**Table S5. Pangenome Composition of Different *B. longum* Subspecies.**

**Table S6. The Proportion of *B. longum* Subspecies in Westernized and Non-Westernized Infants.**

**Table S7. Summary of the metagenome data used in this study.**

**Table S8. Summary of the cattle metagenome data used in this study.**

**Table S9. Summary of the non-human primate's metagenome data used in this study.**

**Table S10. Summary of the ancient human's metagenome data used in this study.**

**Table S11. The publicly available high-quality *B. longum* genomes in NCBI.**

**Table S12. The Published Representative Genomes of Key HMO-metabolizing Strains in infants.**
